# Genome-wide association study and genomic prediction of lucerne traits shaping living mulch performance

**DOI:** 10.64898/2026.04.28.721352

**Authors:** Zineb El Ghazzal, Marie Pegard, Martin Guacaneme, Fabien Surault, Irving Arcia-Ruiz, Gaëtan Louarn, Bernadette Julier

## Abstract

Lucerne is gaining interest as a living mulch in agroecological productions. However, its vigorous growth can lead to competition with cash crops for light and nutrients, necessitating new ideotypes. This study investigated the genetic basis of traits relevant to ideotype breeding: dormancy, spring regrowth, height, growth habit, leaflet size, stem diameter, and plant structure. Individuals from a diversity panel of 27 accessions and a synthetic population were phenotyped in a spaced plant nursery. Over 100,000 SNP markers were used for genotyping. Genome-wide association study (GWAS) and genomic prediction were conducted, considering population structure. Heritability estimates ranged from moderate to high in diversity panel (h² = 0.36–0.70) but were lower in synthetic population (h² = 0.17–0.33), reflecting reduced genetic variance. Trait correlations differed markedly between populations, indicating the possibility of recombining traits to create new ideotypes. GWAS identified a few QTL (r² up to 0.27) for leaflet size, height, growth habit, and plant structure, with candidate genes linked to growth, stress response, and signalling pathways. Genomic prediction was highly accurate in diversity panel, where broad genetic variation allowed reliable estimation of marker effects, with prediction accuracies exceeding 0.8 for heritable traits, including growth habit and leaflet size. In contrast, accuracies were low in synthetic population, reflecting its limited diversity and small size, whether training was based on the synthetic population itself or on the diversity panel. These results highlight the potential to recombine traits and develop lucerne ideotypes using molecular tools such as QTL detection and genomic prediction.

## 1. Introduction

Lucerne (*Medicago sativa* L.) is often referred to as the "queen of forages" because of its high yield per unit area and exceptional nutritional quality. It is one of the most widely cultivated forage legumes worldwide, thanks to its adaptability to diverse climatic and soil conditions (Julier et al. 2017). As a perennial legume, lucerne provides long-term agronomic and ecological benefits. Its nitrogen-fixing capacity improves soil nitrogen content, while its deep root system accesses to subsoil moisture and enhances physicochemical soil properties (Rashmi et al. 1997; Song et al. 2019; Tucak et al. 2021). Furthermore, its dense canopy competes with weeds, providing significant weed suppression in the rotations (Meiss et al. 2010; Barilli et al. 2017). However, with current decoupling of plant and animal productions at the geographic scale, some agricultural landscapes are now composed of annual crops only (Garrett et al. 2020). Lucerne is therefore gaining interest as a service plant in agroecological productions, particularly when used as a living mulch intercropped with cash crops (Cougnon et al. 2022).

Living mulches, defined as cover crops sown prior to the cash crop, persist as a living ground cover throughout and beyond the crop growing season (Hartwig and Ammon 2002; Carof et al. 2007a; Leoni et al. 2022). To maintain their role as service plants, living mulches must ensure rapid and dense ground coverage for effective weed suppression (Leoni et al. 2020) and contribute to nitrogen enrichment, as biomass accumulation is closely linked to nitrogen fixation (Louarn et al. 2015). Perennial legumes, such as lucerne, are the most common species used as living mulches in cereal systems, as their ability to fix atmospheric nitrogen improves soil fertility and reduces the need for fertilisers (Cougnon et al. 2022). They can also support the reduction of intensive tillage practices, thereby saving energy and preserving biodiversity (Vincent-Caboud et al. 2019; Fonteyne et al. 2020). Despite its potential benefits, lucerne faces several challenges when used as a living mulch. Current varieties, which have been selected for forage production, exhibit vigorous growth, leading to significant competition with cash crops for key resources, particularly light and nutrients (Carof et al. 2007b; Cougnon et al. 2022; El Ghazzal et al. 2025). This competition often results in yield losses for the cash crop; for example, wheat yield has been shown to decrease by an average of 45% (Humphries et al., 2004), reaching up to 50% in some cases (Shili-Touzi 2009).

Different strategies can be employed to manage the competitive effects of lucerne used as a living mulch. While mechanical and chemical methods are available, they often raise concerns regarding cost, labour, and environmental impact (Cougnon et al. 2022). A promising alternative involves the selection of less competitive lucerne varieties. Defining an effective ideotype requires balancing two contradictory objectives: minimising competition with the cash crop while maintaining enough competitiveness to suppress weeds. Several key traits must be considered to optimise lucerne growth and compatibility with cash crop, by enhancing its ability to suppress weeds during the off-season or early development of the cash crop, while avoiding light competition during the cash crop growth cycle (Carof et al. 2007b). An appropriate ideotype would combine late spring regrowth, high autumn dormancy, and slow spring regrowth to limit competition (Cougnon et al. 2022). This aligns with findings by Humphries et al. (2004), reporting reduced wheat yield losses when using winter-dormant lucerne varieties (dormancy classes 0.5 and 2) compared with winter-active ones (classes 6 and 10). Moreover, El Ghazzal et al. (2025) showed that varieties combining intermediate dormancy, moderate height, slow spring regrowth, and high lodging resistance would provide the best balance between ecological services, such as weed suppression, and compatibility with the cash crop.

A better understanding of the genetic basis of key traits is essential to accelerate breeding efforts and effectively develop lucerne ideotypes adapted for living mulch use. Lucerne exhibits extensive phenotypic and genetic diversity, shaped by multiple biological and ecological factors, including both natural and artificial selection. While domestication and selection have narrowed the genetic base used in cultivated populations (Muller et al. 2006), wild accessions retain broad variation, reflecting adaptation to climates ranging from cold to arid (Julier et al. 1995). Complex population structure and significant relatedness among accessions have been observed (Li and Brummer 2009; Pégard et al. 2023). This diversity makes the *Medicago sativa* complex, comprising mainly the *sativa* and *falcata* subspecies, a valuable reservoir for breeding programmes targeting sustainable agricultural practices, particularly ideotypes for living mulch systems (Cougnon et al. 2022). In addition, as an autotetraploid (2n = 4x = 32) and an outcrossing (allogamous) species, lucerne maintains high levels of heterozygosity and genetic variation within a population or variety, thus hosting a large source of allele diversity (Julier et al. 2000; Musial et al. 2002; Flajoulot et al. 2005; Alfaifi et al. 2013).

Genome-wide association studies (GWAS) and genomic selection (GS) are proposed to exploit lucerne diversity and identify the genetic basis of complex traits (Pégard et al. 2023). The genetic structure within a population can introduce confounding effects in GWAS, resulting in the identification of spurious associations (false positives) due to ancestry differences and cryptic relatedness (Kaler et al., 2020; Sul et al., 2018; Thornton et al., 2012). To address this issue, mixed linear model (MLM) based GWAS methods have been widely adopted in structured plant populations, including the multi-locus mixed model (MLMM) (Segura et al. 2012). These models are effective at controlling false positives by considering relatedness with a kinship matrix and sequentially incorporating associated markers as cofactors through forward and backward selection (Alemu et al. 2021; Alamin et al. 2022). Genomic selection, on the other hand, is increasingly used in plant breeding programmes to predict the genomic estimated breeding values (GEBV) of individuals based on the cumulative effects of genome-wide markers. Its effectiveness lies in capturing both large- and small-effect loci across the genome, including those often missed in GWAS (Crossa et al. 2017). Prediction accuracy depends on several factors, such as marker density, training population size, trait heritability, and, critically, the genetic relatedness between training and target populations (Pégard et al. 2020). Accuracy generally increases when these factors are optimised. In lucerne, genomic selection has shown moderate to high predictive abilities for complex traits (Li et al. 2015; Annicchiarico et al. 2015b; Pégard et al. 2023), which is encouraging given the similarly complex nature of the traits of interest for living mulch.

GWAS and GS studies in lucerne have employed a range of population designs, each with specific strengths and limitations in terms of allelic diversity, mapping resolution, and prediction accuracy. Diversity panels composed of landraces, cultivars, and wild accessions capture broad allelic variation and enable high-resolution mapping, although they require careful correction for population structure (Pégard et al. 2023, 2025). Breeding panels, while genetically narrower, are highly relevant to applied breeding and often yield moderate to high prediction accuracies due to close relatedness between training and target populations (Li et al. 2015). F₁ full-sib or half-sib populations allow precise segregation analyses and detection of rare alleles, though they capture only a limited fraction of diversity (Murad Leite Andrade et al. 2022). Synthetic populations derived from multiple parents or commercial cultivars offer intermediate diversity and reduced structure, which can increase the power to detect true associations in GWAS (Biazzi et al. 2017).

In this study, two populations, a diversity panel and a synthetic population, were analysed to investigate the genetic basis of traits relevant to lucerne ideotype breeding for living mulch systems. Specifically, autumn dormancy, spring regrowth rate, plant height, growth habit, leaflet size, stem diameter, and plant structure were studied. Both populations were phenotyped and genotyped using high-density SNP markers, and trait heritability and genetic correlations were estimated. GWAS was conducted to identify associated loci, and genomic prediction models were tested within and across populations to evaluate the potential of GS for creating lucerne ideotypes. The study also assessed how prediction accuracy depends on heritability and genetic relatedness, providing insights for designing effective breeding strategies for lucerne varieties suited for living mulch use.

## 2. Materials and Methods

### 2.1. Plant material

Two populations, differing for composition and diversity range, were used. The “accession panel” was composed of 27 tetraploid *Medicago sativa* accessions selected for their high morphological variability, ranging from wild prostrate to tall, erect cultivated types, each represented by 40 individuals (Table S1). These accessions belonged to two subspecies within the *M. sativa* complex: subsp. *sativa* and subsp. *falcata*. The synthetic population “Syn1”, was composed of 228 individuals of the first-generation progeny of a polycross between three populations, Krasnokutskaya (subsp. *falcata*), Mezzo (subsp. *sativa*), and LPIII (subsp. *sativa*). The first two were part of the accession panel, and LPIII was included for its high seed production. Each parental population was represented by 30 genotypes in the polycross. The accession panel is supposed to cover a wider range of diversity and to show larger genetic structure than the Syn1.

### 2.2. Experimental design

Seeds were scarified to enhance germination, then incubated at 25 °C in the dark in 10 cm Petri dishes lined with moist filter paper. After germination, seedlings were transplanted into plastic pots and grown in a greenhouse for one month. For each accession of the panel, 40 healthy plants were transplanted into the field on 21 April 2021. The same procedure was applied to the plants from the Syn1 polycross progeny. The field received an initial irrigation of approximately 25 mm following transplanting, and then relied solely on natural rainfall (Table S2). The experiment was conducted from April 2021 to June 2023 at the INRAE research station in Lusignan, France (46°26′08″N, 0°07′25″E), on clay loam soil with an acidic pH of 6.5. The field was organised into four blocks. Within each block, 10 plants per accession of the diversity panel were planted in 10 columns spaced 0.7 m apart, with each row corresponding to a distinct accession. The space between rows was 0.7 m too. The accession "Milky Max" was included in four rows in each block in different spatial positions to assess spatial variability. The Syn1 was planted in the same field, at the end of each block after the accession rows (Figure S1). Weed control was performed mechanically between plants, with repeated passes as needed throughout the duration of the trial.

### 2.3. Genotyping and SNP filtering

Leaflet sampling, DNA extraction, and GBS library construction were performed at the plant level, following the method outlined by Julier et al. (2018), with a modified digestion protocol involving a double digestion using the restriction enzymes *PstI* and *MseI*. SNP calling and allele frequency determination followed the pipeline described by Pégard et al. (2023). Briefly, raw reads were processed using the GBprocesS pipeline, which included demultiplexing, trimming, merging, quality filtering, and mapping to the haploid *Medicago sativa* ‘B’ reference genome (Chen et al. 2020). Genotyping was performed by extracting read counts for each nucleotide position, retaining only those with a read depth between 10 and 1200. Only biallelic SNP with a minor allele frequency (MAF) above 1% were selected.

Quality filtering was conducted separately for the accession panel and the Syn1 to account for differences in allele frequencies. For both datasets, individuals with more than 60% missing data were removed, as were SNP with more than 20% missing values or with MAF below 5%. After filtering, 111,486 SNP were retained for the accession panel, including 42,894 without missing data and an overall missing rate of 2.7%. For the Syn1, 106,750 SNP were retained, among which 57,040 showed no missing data, with a missing rate of 2.4%. Missing genotype data were imputed separately for the accession panel and the Syn1, using the mean allele frequency of each SNP. For the accessions, imputation was performed at the accession level: missing genotypes for a given individual were imputed using the mean allele frequency of that SNP among other individuals from the same accession.

### 2.4. Genetic diversity

To characterise the genetic structure of the accession panel, a principal component analysis (PCA) was first performed using the R package “*FactoMineR”* (Lê et al. 2008), based on 39,400 SNP common to both the accession panel and the Syn1, without missing data. Principal components accounting for 80% of the total genetic variation were retained for subsequent clustering. Syn1 individuals were then projected onto the same PCA as supplementary individuals to determine their genetic position relative to the accession panel. Subsequently, a discriminant analysis of principal components (DAPC) was conducted using the R package “*adegenet”* (Jombart and Ahmed 2011). The optimal number of genetic clusters was identified using the “*find.clusters*” function, based on the Bayesian information criterion (BIC), by selecting the first local minimum of the BIC. Genetic differentiation among clusters was estimated using pairwise F_ST_ (Weir and Cockerham 1984). Linkage disequilibrium (LD) decay was assessed by calculating the squared partial correlation coefficient (r²) for each intrachromosomal SNP pair (Mangin et al. 2012; Lin et al. 2012), to evaluate the extent of LD in the panel and the Syn1 and to estimate the expected resolution of association mapping.

### 2.5. Phenotypic Data

Autumn regrowth was used as a proxy for dormancy, where lower regrowth indicates higher dormancy, following the ‘Fall Dormancy’ standard test (Teuber et al. 1998). After the summer harvest on 16/09/2021, plant height was measured three times from the soil to the top of the tallest stretched stem. These three measurements were used to calculate the slope of autumn regrowth over time, expressed in cumulative thermal time (growing degree days (GDD), base 0 °C, in cm ·GDD⁻¹. Spring regrowth was characterised by three traits. First, the regrowth rate was calculated as the slope of stretched height (cm·GDD⁻¹) recorded every 15 days from March to May 2022. Second, the last measurement in May was taken as the maximum height, and finally, leaflet size was visually assessed using a scale from 1 (small) to 9 (large). Growth habit was also visually assessed in June and August 2021, with ratings ranging from 1 (upright growth) to 9 (prostrate growth), before any occurrence of lodging. Plant structure was described by the ratio of maximum natural height to maximum stretched height, both measured on 13/05/2022. This index reflects not only lodging but also the influence of growth habit. In prostrate accessions, which develop close to the soil surface, the combination of a naturally short standing height and a much greater stretched height may yield values similar to lodged plants, although these accessions are not truly lodged. Stem diameter (mm) was calculated as the average diameter of three randomly selected stems per plant, measured at the lower third of the stem in May 2022 and 2023. This trait was included as a key determinant of lodging resistance, reflecting the mechanical support provided by stem tissues (Guines et al. 2003).

To correct the microenvironmental heterogeneity within the experimental trial, phenotypic data for each trait were independently adjusted for spatial variability using R package “breedR” (Munoz and Rodriguez 2020). The adjustment was carried out using a linear mixed model (Eq. 1), that incorporated both genetic effects, spatial effects across rows, columns, and blocks.

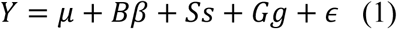

Where *Y* was the vector of phenotypic observations, and ***μ*** was the overall mean. *Bβ* represented the fixed effects of blocks, where *B* was the incidence matrix for blocks and *β* was the vector of block effects. *Ss* represented the random spatial effect across rows and columns within the field, following *s* ∼ ℕ(0, *Sσ*_*s*_²), where *S* was the covariance matrix derived from a two-dimensional tensor product of B-splines along rows and columns, and *σ*_*s*_² was the spatial variance. *Gg* represented the random additive genetic effects, g ∼ ℕ (0, G*σ*_*g*_²), where *G* was the Genomic Relationship Matrix (GRM), and *σ*_*g*_² was the additive genetic variance. Finally, *ϵ* represented the residual error, where *ϵ* ∼ ℕ(0, *Iσ*_*e*_²), with *I* was the identity matrix and *σ*_*e*_² was the residual variance. The GRM was computed following (Eq.2).

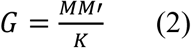

Where *M* was the *n* × *q* matrix (*n*: number of individuals, *q*: number of SNP), centred by allele frequency following VanRaden (2008) method. Only SNP without missing values were used to compute *M*. The matrix was scaled so that the average of the diagonal elements equalled 1, following Forni et al. (2011) (Eq.3).

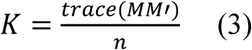

The same mixed model (Eq. 1) described above was used to estimate narrow sense heritability (h²) of each trait (Eq. 4), as well as the genetic correlation between traits using the adjusted phenotypic data (Eq. 5).

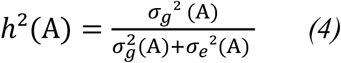

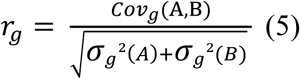

where *Cov*_*g*_(*A*, *B*) was the genetic covariance between trait A and trait B. Phenotypic correlation among traits was estimated on adjusted phenotypic data, using Pearson coefficient.

The effect of genetic group on phenotypic variation was assessed using analysis of variance (ANOVA). When significant differences among groups were detected, Tukey’s Honest Significant Difference (HSD) test was applied to perform pairwise comparisons between genetic groups.

### 2.6. GWAS and QTL Analysis

GWAS was conducted separately for the accession panel (111,486 SNP) and the Syn1 (106,750 SNP). Analyses were performed using the MLMM approach (Segura et al. 2012), accounting for population structure and genetic relatedness, and implemented with the R package “*mlmm*”. Each trait was analysed using stepwise mixed-model regression. Significant associations were identified based on a Bonferroni-adjusted threshold (0.1 / number of SNP). The proportion of phenotypic variance explained (*r*²) by each QTL was estimated by comparing mixed linear models with and without the focal QTL using likelihood ratio tests. A combined *r*² was also calculated by including all significant QTL. Finally, candidate genes located near significant QTL were identified using the *Medicago sativa* reference genome (Chen et al., 2020) through the “*myGenomeBrowser*” platform (Carrere and Gouzy 2017).

### 2.7. Genomic prediction

To evaluate the predictive ability of genomic selection, four scenarios (S1–S4) were tested using the Genomic Best Linear Unbiased Prediction (GBLUP) model, implemented in the R package “*breedR*” (Muñoz & Rodríguez, 2020). The first two scenarios assessed within-population prediction. In Scenario 1 (S1), prediction was performed within the accession panel (n = 1025) using 111,486 SNP and stratified cross-validation. A fixed 20% (n = 205) of individuals was randomly selected as a test set, while the remaining 80% was split into training subsets ranging from 20% to 80% of the population and validation subsets. Stratification across genetic clusters was applied to preserve population structure, as this improves predictive ability when genetic subgroups are represented in the training set (Pégard et al. 2023). This scenario simulated selection within structured germplasm and evaluated prediction performance across a genetically diverse panel of accessions. Scenario 2 (S2) assessed prediction within the Syn1 population (n = 228) using 106,750 SNP. Here, 30% of individuals (n = 68) were used as a fixed test set, while the remaining 70% formed training subsets of increasing size (20% to 80% of Syn1 plants), maintaining balanced representation from the three Syn1 parental accessions. This scenario reflects early-stage selection in a synthetic breeding pool and enables the identification of superior genotypes within targeted diversity.

The last two scenarios addressed across-population prediction. Scenario 3 (S3) used the full accession panel as the training population, with the Syn1 as the test population, based on the 104,250 SNP common to both datasets. Stratified sampling was again applied to training subsets (20% to 80%) to retain population structure. This scenario examines the feasibility of predicting phenotypes in material with a reduced diversity using a very diverse germplasm panel, which is relevant for pre-breeding to reduce phenotyping costs. Scenario 4 (S4) followed the same structure as S3 but limited the training population to cultivated accessions from three genetic clusters, including two of the Syn1 parents. This design represents a more realistic breeding scenario with moderate genetic relatedness between training and test populations and helps assess the impact of relatedness on prediction ability. All scenarios were replicated 30 times to ensure robust estimation. Predictive ability was evaluated as the Pearson correlation between genomic estimated breeding values (GEBV) and adjusted phenotypes.

## 3. Results

### 3.1. Population Structure and Genetic Relatedness

In the PCA conducted to assess the genetic structure of the accession panel, the first two principal components explained 10.4% (PC1) and 2.2% (PC2) of the total genetic variation. Clustering analysis of the accessions identified six distinct genetic groups (K = 6; Figure S2), which broadly aligned with subspecies and cultivation status (Figure 1A). Groups G1 and G2 comprised cultivated *M. sativa* subsp. *sativa* accessions, although individuals were dispersed between these two clusters. Group G3 comprised non-dormant cultivated *sativa* types, including the Tunisian landrace ‘Gabès’ and the variety ‘Speeda’. Group G4 contained three wild *sativa* accessions from Spain (‘Mielga’), the turf-type variety ‘GreenMed’ (derived from ‘Mielga’), and a Chinese cultivated accession (‘Wugong’). Group G5 consisted of cultivated *falcata*, while G6 included wild *falcata* accessions. When the Syn1 individuals were projected onto the PCA (Figure 1A), they occupied an intermediate position between cultivated *sativa* (G1, G2, G3) and *falcata* (G5), reflecting their mixed background.

**Figure 1.**
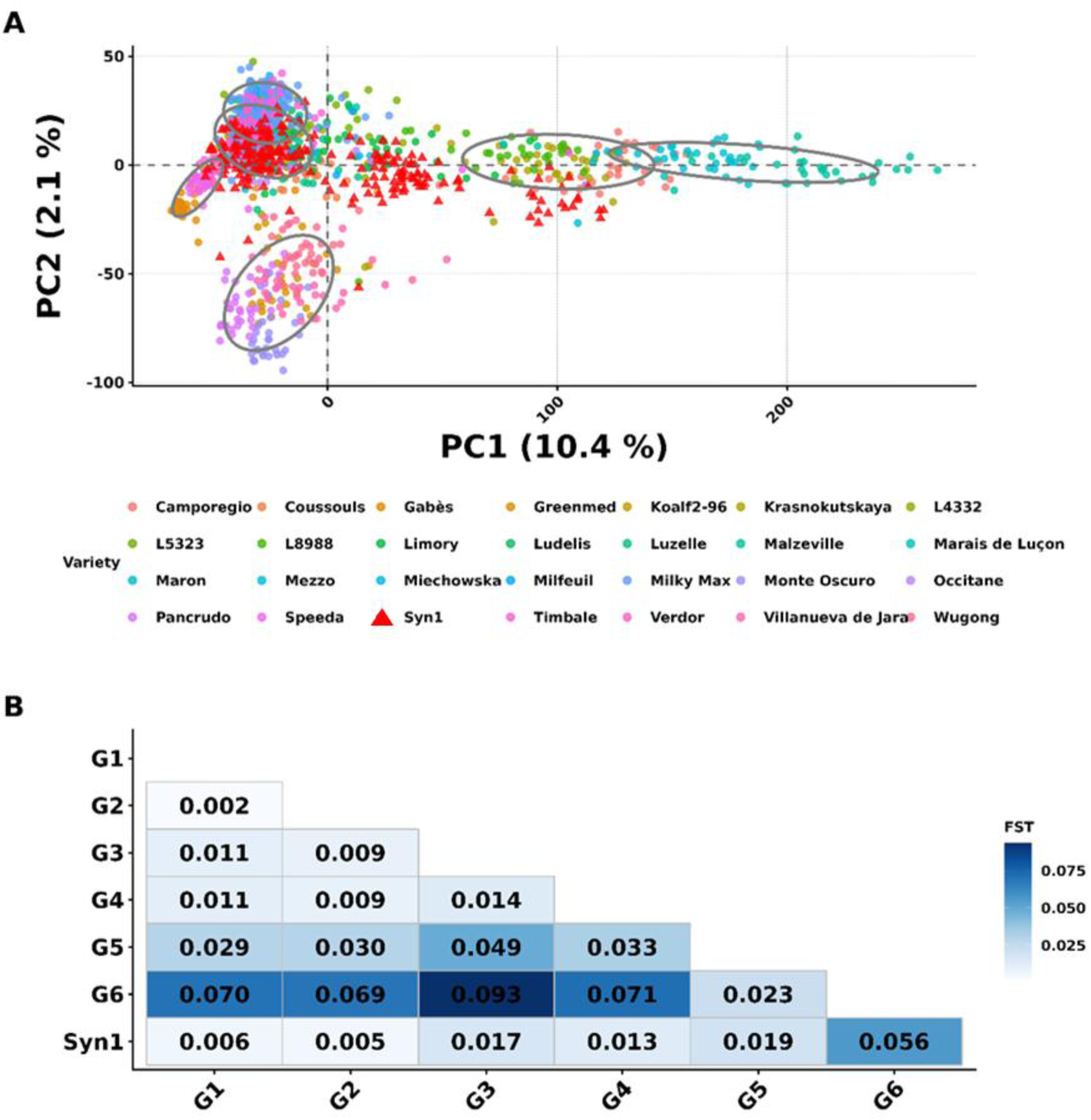
Genetic structure and linkage disequilibrium in the lucerne accession panel and the Syn1 population. (A) Principal component analysis (PCA) of individual plants from the 27-accession panel with projection of the Syn1 individuals, based on 39,400 SNP. Circles indicate cluster assignments (G1–G6), and ellipses represent 80% confidence envelopes for each cluster. (B) Pairwise genetic differentiation (FST) among the six clusters and between each cluster and the Syn1.

Pairwise F_ST_ estimates among the six clusters revealed a consistent pattern of increasing genetic differentiation from cultivated to wild types, and from *sativa* to *falcata*. F_ST_ values ranged from 0.002 between G1 and G2 (both cultivated *sativa*) to 0.093 between G6 (wild *falcata*) and G3 (non-dormant cultivated *sativa*) (Figure 1B). The Syn1 also showed differentiation from all clusters, with F_ST_ values ranging from 0.005 with G1 (which shared a *sativa* parent) to 0.056 with G6 (*falcata*).

### 3.2. Phenotypic variation

Phenotypic traits varied across both the accession panel and the Syn1 population, with significant differences observed among the six genetic clusters (G1–G6) (Figure 2, Table S3). Dormancy, expressed as autumn growth rate, was more intense in the accession panel (0.037 ± 0.016 cm GDD⁻¹) than in the Syn1 (0.041 ± 0.011 cm GDD⁻¹). In contrast, Growth habit was slightly higher in the Syn1 (4.90 ± 1.00) than in the accession panel (4.65 ± 1.97). Leaflet size followed a similar pattern, being higher in the Syn1 (5.65 ± 0.79) than in the accession panel (5.30 ± 1.34). Spring regrowth rate showed only marginal differences between groups (0.102 ± 0.021 vs. 0.099 ± 0.023 cm GDD⁻¹). Likewise, maximum height and stem diameter were slightly higher in the Syn1 (78.4 ± 14.1 cm vs. 74.2 ± 17.0 cm, and 2.76 ± 0.77 mm vs. 2.45 ± 0.95 mm, respectively). Plant structure scores also tended to be higher in the Syn1 (0.81 ± 0.11) than in the accession panel (0.66 ± 0.18). Although outlier values extended to similar ranges in both groups (Table S3), the diversity in the Syn1 was generally narrower than in the panel (Figure 2).

**Figure 2.**
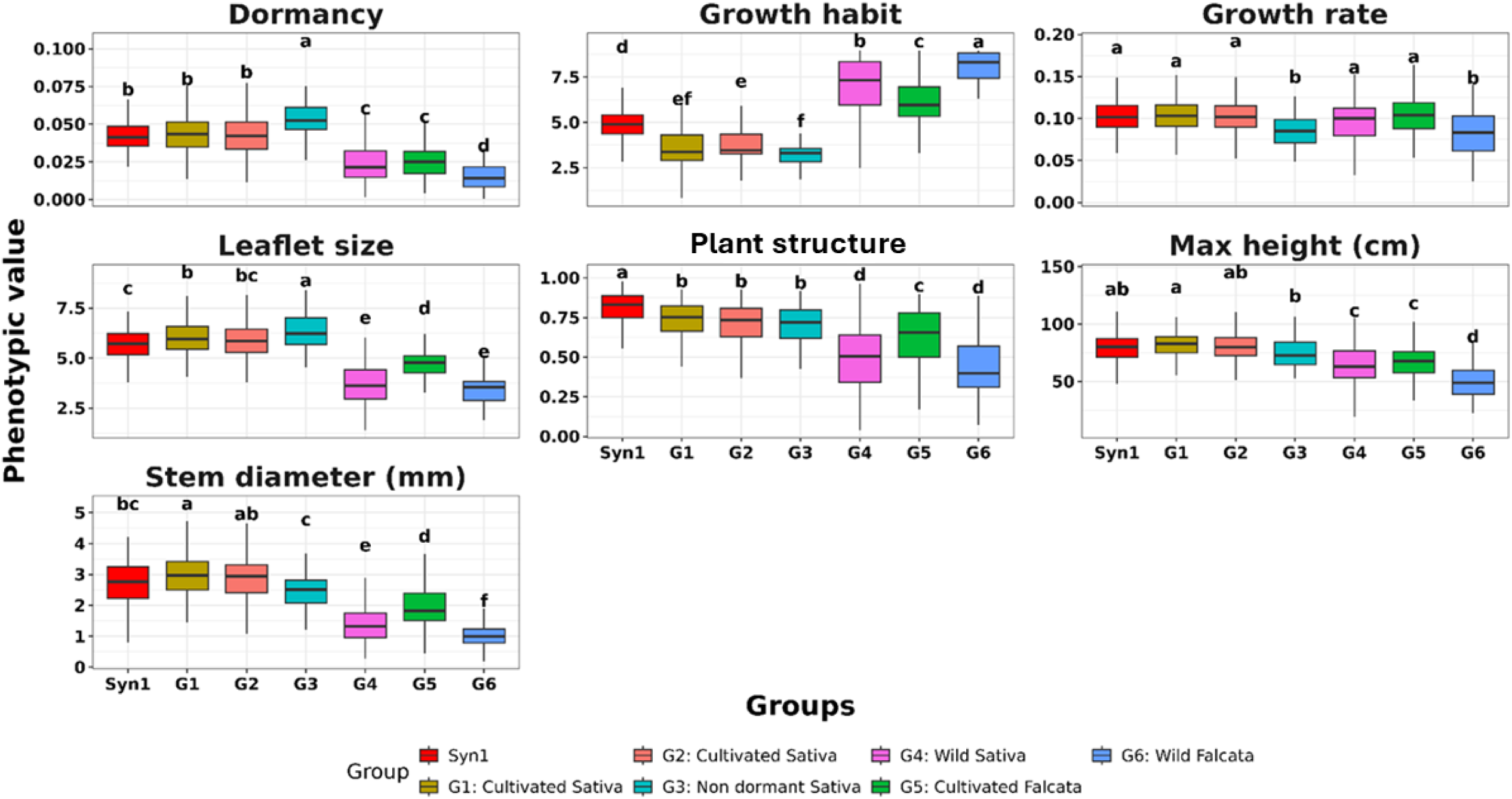
Boxplots illustrating the distribution of phenotypic traits across the six genetic clusters (G1–G6) of the accession panel and the Syn1 population. The letters above the boxplots denote the statistically significant differences among groups based on Tukey’s HSD test (p < 0.05) following ANOVA.

Analysis of variance (ANOVA) confirmed that the genetic cluster had a significant effect on all traits (p < 0.001), with pairwise differences identified via Tukey’s HSD test (Figure 2). Cultivated *sativa* accessions (G1–G3) were characterised by great height, large leaflets, fast regrowth, thick stems, and good Plant structure. They also exhibited erect architecture (low growth habit scores) and intermediate dormancy. Among them, G3 displayed the lowest autumn growth rate, corresponding to the highest dormancy. In contrast, G5 (cultivated *falcata*) exhibited intermediate values, and G4 and G6 (wild *sativa* and *falcata*, respectively) showed the lowest trait values overall for dormancy. The Syn1 population broadly showed a similar range of variation as the cultivated *sativa* groups, but its variation did not cover that included in the wild accessions of the diversity panel.

Trait heritability estimates (Table S3) ranged from moderate to high in the accession panel (h² from 0.36 for spring growth rate to 0.70 for growth habit). In contrast, the Syn1 showed lower heritability across all traits (h² = 0.17 to 0.33), largely reflecting reduced genetic variance, while strong spatial or residual effects also contributed for some traits.

### 3.3. Genetic and phenotypic correlations

Genetic and phenotypic correlations among traits were estimated separately for the accession panel and the Syn1 population (Figure 3). In the accession panel (Figure 3A), growth habit and leaflet size were strongly negatively correlated (genetic *r* = –0.62; phenotypic *r* = –0.73), indicating that prostrate genotypes tended to have small leaflets. Dormancy (as slow autumn growth) was positively correlated with leaflet size (genetic *r* = 0.60; phenotypic *r* = 0.52), while spring regrowth rate and maximum height were highly positively correlated (genetic *r* = 0.78; phenotypic *r* = 0.79). In contrast, stem diameter exhibited near-zero genetic correlation with height (*r* = –0.05) but a strong positive phenotypic correlation (*r* = 0.69), and also showed strong phenotypic correlations with leaflet size (*r* = 0.70) and growth habit (*r* = -0.67). Plant structure was moderately negatively correlated with growth habit (genetic *r* = –0.47; phenotypic *r* = –0.56), indicating that more erect genotypes were better able to resist lodging.

**Figure 3.**
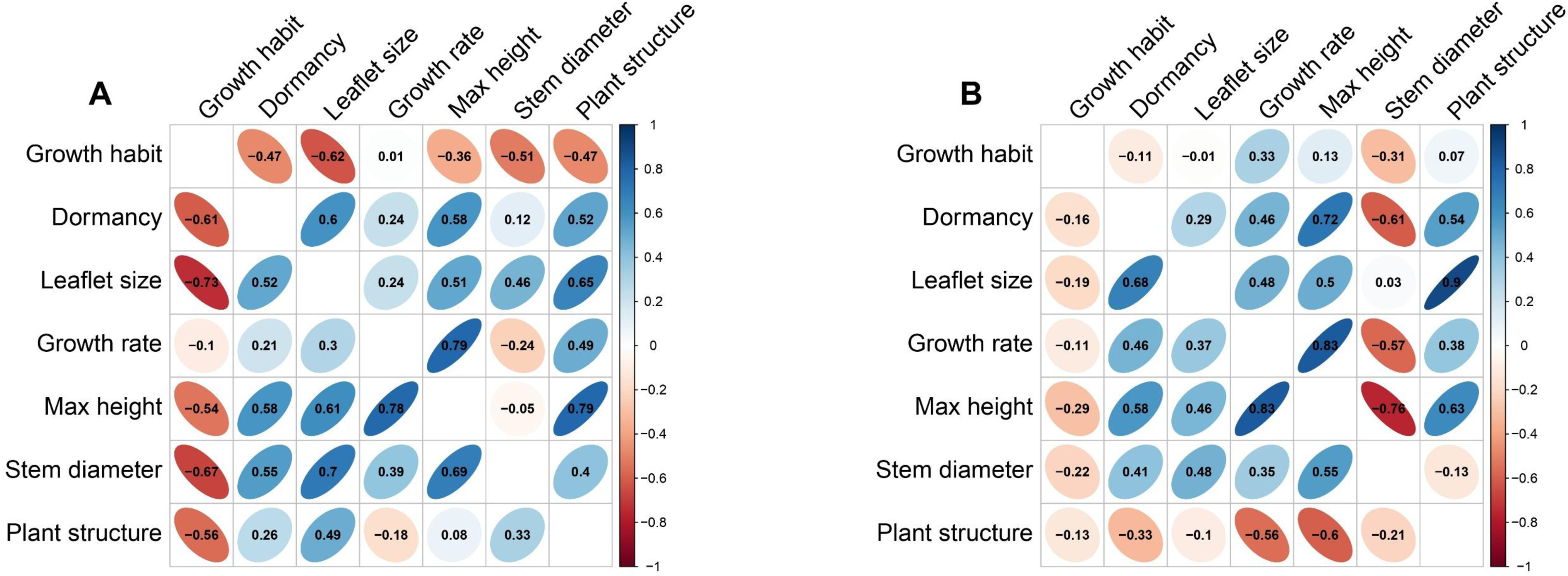
Heatmaps of genetic and phenotypic correlation coefficients between traits, with the upper triangle representing genetic correlations and the lower triangle showing phenotypic correlations, in the accession panel (A) and the Syn1 population (B). (p < 0.05 (*), p < 0.01 (**); p < 0.001 shown without asterisks)

In the Syn1, the correlation matrix was very different. Many genetic correlations were reduced or reversed relative to the accession panel, and the alignment between genetic and phenotypic correlations was less consistent (Figure 3B). The strong negative phenotypic and genetic correlation between growth habit and leaflet size observed in the accessions was reduced to a weak correlation in the Syn1 (r = –0.19 and r = –0.01, respectively). More generally, all correlations involving growth habit were weak in the Syn1, consistent with the reduced variability of this trait compared to the accession panel. The dormancy–leaflet size correlation also declined to r = 0.29 genetically and r = 0.11 phenotypically. Stem diameter showed a strong negative genetic correlation with dormancy (r = –0.61). It was also strongly negatively correlated with height at the genetic level (r = –0.76), in contrast with the positive phenotypic correlation between stem diameter and height (r = 0.55), which reflected the expected structural relationship between these traits. Plant structure, which had been largely independent of height in the accession panel, became strongly positively correlated (genetic *r* = 0.63; phenotypic *r* = 0.60). The correlation between growth habit and plant structure also shifted from strongly negative in the panel to near-zero in the Syn1 (genetic *r* = 0.07; phenotypic *r* = –0.13).

### 3.4. Genome-wide association studies (GWAS)

Linkage disequilibrium (LD) decayed rapidly in both populations, but at different scales. In the accession panel, mean r² dropped below 0.1 within approximately 200 base pairs across all chromosomes (Figure S3A), with over 99.95% of SNP pairs showing r² values below 0.1. In contrast, LD decay was slower in the Syn1, with mean r² falling below 0.2 at ∼1000 bp (Figure S3B). Based on these patterns, candidate gene search windows were set to ±2 kb in the accession panel and ±5 kb in the Syn1.

In the accession panel, a major QTL on chromosome 7 was associated with both growth habit and leaflet size, explaining 27.3% and 27.6% of phenotypic variance, respectively (Table 1, Figure S4). This locus lies 795 bp upstream of MS.gene036920, a RING-type zinc finger protein involved in protein ubiquitination. A minor QTL for spring regrowth rate was identified on chromosome 8 (3.8% of phenotypic variance), located near MS.gene61257, a calcium-binding protein. Plant structure was associated with a QTL on chromosome 6 (15.6% of phenotypic variance), near MS.gene031124, encoding a ubiquitin conjugation factor E4. In Syn1, three QTL for leaflet size were detected on chromosomes 1, 2, and 4, jointly explaining 31% of phenotypic variance. Only the QTL on chromosome 4 was near a gene (MS.gene91452), though its function was not annotated. For plant height, multiple QTL were found on chromosomes 1, 2, 3, 4, 5, and 8, cumulatively explaining 60% of phenotypic variance. A major QTL on chromosome 8 (30% of phenotypic variance) was located in MS.gene000456, encoding a protein involved in ubiquitin signalling. Other loci were associated with defence, chromatin organisation, and growth pathways (Table 1, Figure S5). Overall, GWAS revealed distinct QTL architectures between the two populations, with no overlapping loci identified.

**Table 1.**
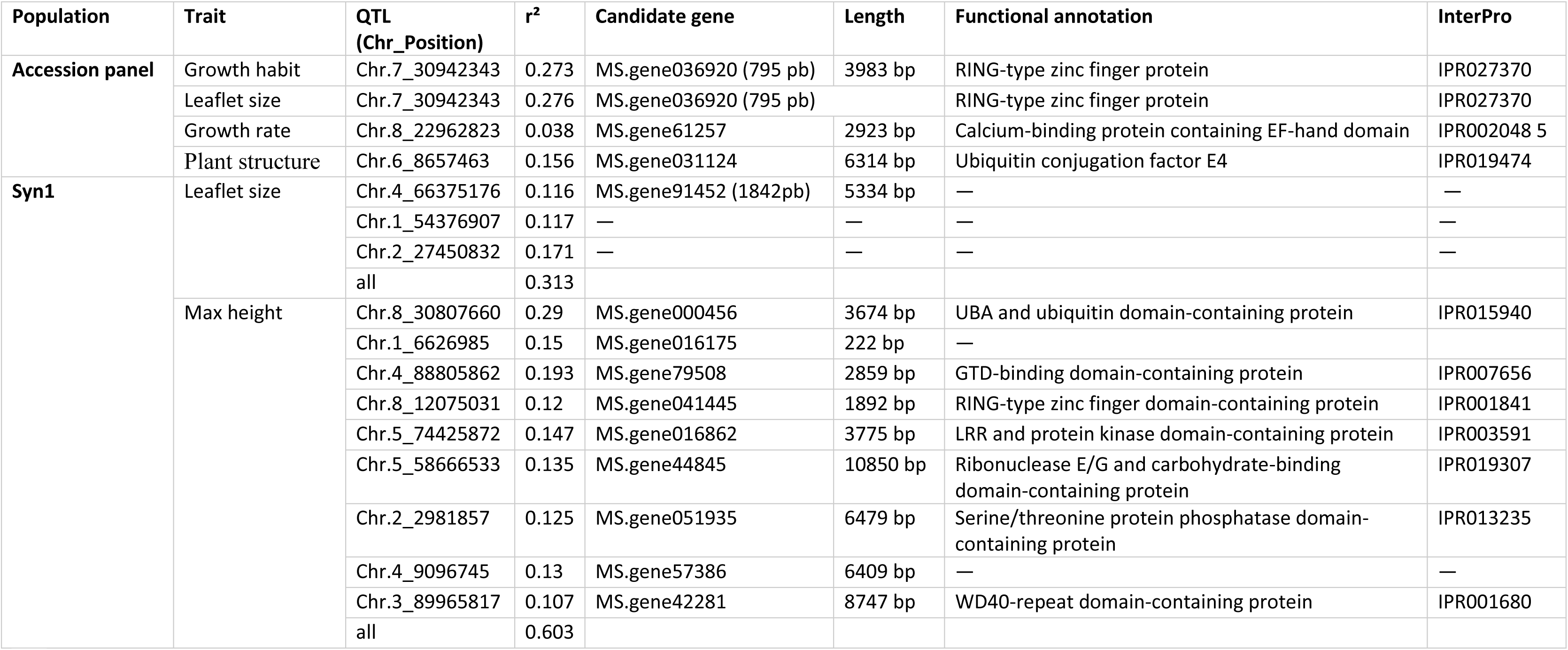
Summary of significant QTLs identified in the accession panel and the Syn1 population. For each QTL, the chromosome and the physical position, as well as the proportion of phenotypic variance explained (r²), are reported. Gene information includes the gene ID (with the distance noted when the QTL is located outside the gene), gene length, predicted functional annotation, and associated InterPro domains.

| Population | Trait | QTL<br>(Chr_Position) | $r^2$ | Candidate gene | Length | Functional annotation | InterPro |
| --- | --- | --- | --- | --- | --- | --- | --- |
| Accession panel | Growth habit | Chr.7_30942343 | 0.273 | MS.gene036920 (795 pb) | 3983 bp | RING-type zinc finger protein | IPR027370 |
|  | Leaflet size | Chr.7_30942343 | 0.276 | MS.gene036920 (795 pb) |  | RING-type zinc finger protein | IPR027370 |
|  | Growth rate | Chr.8_22962823 | 0.038 | MS.gene61257 | 2923 bp | Calcium-binding protein containing EF-hand domain | IPR002048 5 |
|  | Plant structure | Chr.6_8657463 | 0.156 | MS.gene031124 | 6314 bp | Ubiquitin conjugation factor E4 | IPR019474 |
| Syn1 | Leaflet size | Chr.4_66375176 | 0.116 | MS.gene91452 (1842pb) | 5334 bp | — | — |
|  |  | Chr.1_54376907 | 0.117 | — | — | — | — |
|  |  | Chr.2_27450832 | 0.171 | — | — | — | — |
|  |  | all | 0.313 |  |  |  |  |
|  | Max height | Chr.8_30807660 | 0.29 | MS.gene000456 | 3674 bp | UBA and ubiquitin domain-containing protein | IPR015940 |
|  |  | Chr.1_6626985 | 0.15 | MS.gene016175 | 222 bp | — |  |
|  |  | Chr.4_88805862 | 0.193 | MS.gene79508 | 2859 bp | GTD-binding domain-containing protein | IPR007656 |
|  |  | Chr.8_12075031 | 0.12 | MS.gene041445 | 1892 bp | RING-type zinc finger domain-containing protein | IPR001841 |
|  |  | Chr.5_74425872 | 0.147 | MS.gene016862 | 3775 bp | LRR and protein kinase domain-containing protein | IPR003591 |
|  |  | Chr.5_58666533 | 0.135 | MS.gene44845 | 10850 bp | Ribonuclease E/G and carbohydrate-binding domain-containing protein | IPR019307 |
|  |  | Chr.2_2981857 | 0.125 | MS.gene051935 | 6479 bp | Serine/threonine protein phosphatase domain-containing protein | IPR013235 |
|  |  | Chr.4_9096745 | 0.13 | MS.gene57386 | 6409 bp | — | — |
|  |  | Chr.3_89965817 | 0.107 | MS.gene42281 | 8747 bp | WD40-repeat domain-containing protein | IPR001680 |
|  |  | all | 0.603 |  |  |  |  |

### 3.5. Genomic prediction

Prediction ability varied considerably across the four genomic prediction scenarios (S1–S4) (Figure 4A). In S1, conducted within the accession panel, prediction ability (PA) was generally high. Traits such as growth habit and leaflet size achieved PA values above 0.80 even with small training sets, while others, including plant height, stem diameter, and plant structure, reached PA between 0.50 and 0.70 with increasing training set size. Spring regrowth rate exhibited the lowest prediction ability in this panel (PA = 0.33 with the largest training set). S2, conducted within the Syn1 population, showed lower PA than S1. Prediction improved with training set size, but still at moderate levels. The most predictable traits were stem diameter and growth habit (PA = 0.37 at 80%), while dormancy was essentially non-predictable (PA < 0.05), regardless of training set size. Across-population prediction scenarios showed lower accuracy than within-population scenarios. In S3, where the accession panel was used to predict Syn1 performance, PA values were comparable to those observed in S2 for some traits such as growth habit, and only slightly reduced for stem diameter and leaflet size. In contrast, plant structure showed a clear reduction in predictive ability, and PA of other traits that were already poorly predicted in S2 (e.g. dormancy, growth rate) remained low. Restricting the training set to cultivated accessions in S4 produced a similar pattern as S3.

**Figure 4.**
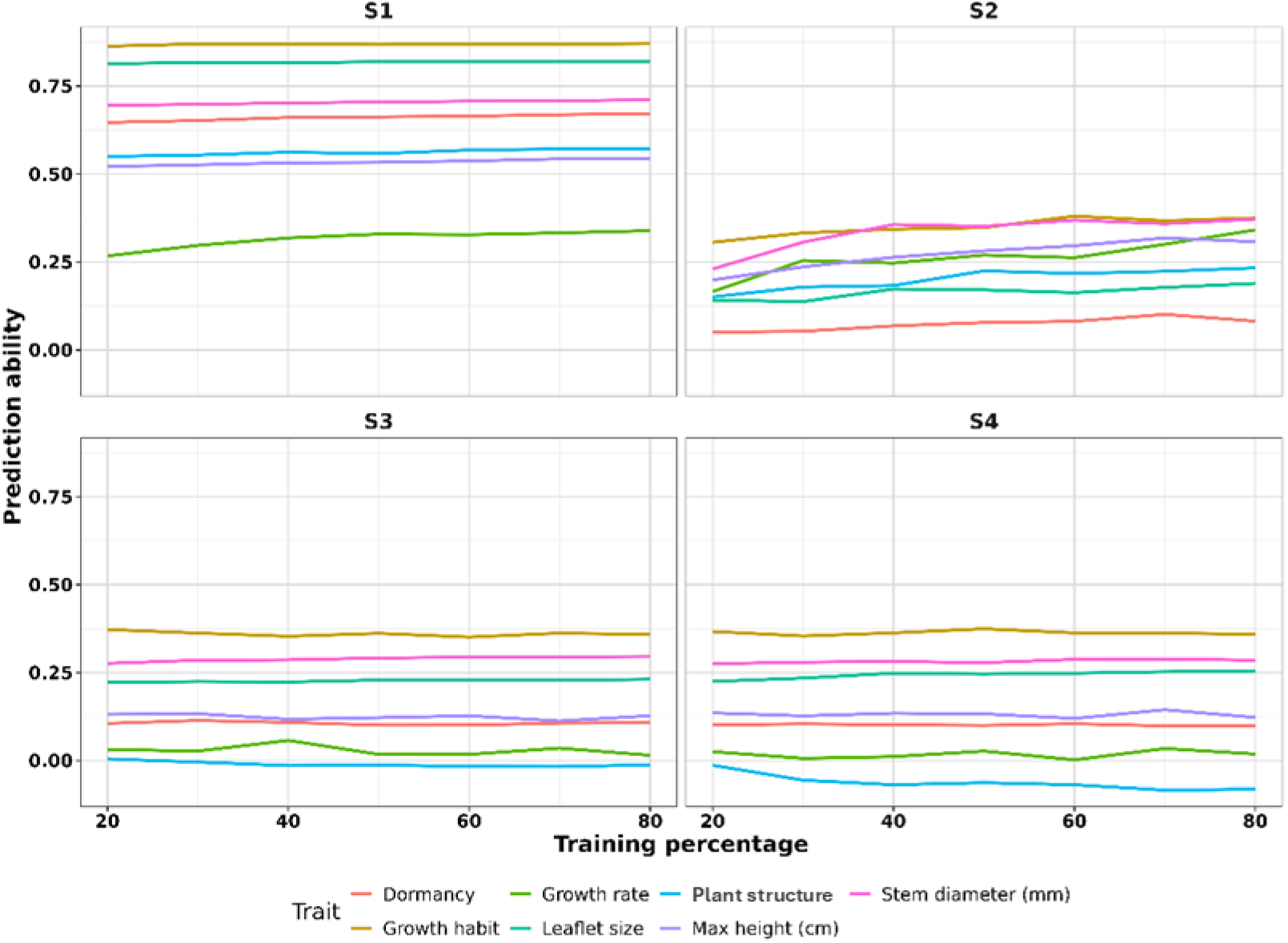
Genomic prediction across scenarios. Prediction ability for the phenotypic traits across four prediction scenarios (S1–S4). evaluated at increasing training set sizes (20% to 80%). Each line represents the mean prediction ability for each trait. In Scenario 1 (S1). predictions were performed within the accession panel. Scenario 2 (S2) prediction was calculated in the Syn1 population. Scenario 3 (S3) tested across-population prediction using the accession panel to predict the Syn1. while Scenario 4 (S4) restricted training to cultivated accessions (G1, G2, G4) of the panel to predict the Syn1.

## 4. Discussion

This study investigated the genetic determinants of lucerne traits linked to dual role in living mulch systems, balancing reduced competition with wheat while maintaining ecosystem services. A diverse panel of accessions and a synthetic population (Syn1) were used to explore genetic variation and assess genomic prediction performance under different population structures.

### 4.1. Phenotypic variation and trait correlations differ between the accession panel and the Syn1

The accession panel exhibited broad phenotypic variation for all traits. Cultivated *M. sativa* subsp*. sativa* groups (G1–G3) were characterised by tall and erect plants with large leaflets, fast regrowth, and thick stems, consistent with traits targeted in forage breeding (Annicchiarico et al. 2015a). In contrast, wild accessions (G4, G6) exhibited prostrate growth, reduced size, and thin stems, reflecting their adaptation to natural environments that allow them to escape grazing. The Syn1 displayed a narrower overall phenotypic range than the accession panel, consistent with its limited number of parents. However, some individuals showed intermediate values, particularly for growth rate and plant structure. These patterns suggest that recombination among divergent parents may have generated unusual phenotypic profiles, even though overall diversity was reduced. These phenotypic patterns aligned with the genetic clusters. The accession panel was structured into six genetic clusters reflecting subspecies and cultivation status, whereas the Syn1 occupied an intermediate genetic position between cultivated *sativa* and *falcata*.

Trait correlation patterns differed substantially between the accession panel and the Syn1, reflecting both the underlying genetic background and the range of diversity captured in each population. In the accession panel, taller plants exhibited stronger plant structure, although height and stem diameter were not genetically correlated. This likely reflects the measurement of maximum stretched height, which captures potential elongation rather than growth habit. Some plants may have long stems yet remain prostrate or semi-erect, whereas truly erect plants tend to develop thick stems to maintain structural support. Leaflet size is a relevant breeding trait due to its influence on canopy structure and photosynthetic efficiency (Simkin 2019). Erect growth habit and large leaflet size were positively associated with plant height, reflecting co-selection for traits promoting light interception and forage yield (Jia et al. 2022).

In the Syn1, correlations between traits were generally weaker than in the accession panel. For example, the strong association between leaflet size and growth habit observed in the accessions was no longer observed, while taller individuals tended to have thinner stems but still resisted lodging. Recombination among divergent parents may have disrupted historical associations between traits, generating novel combinations not present in the accession panel. In addition, this attenuation of correlation values can be partly explained by the narrow range of variation in the Syn1, which limits the detectability of phenotypic relationships.

### 4.2. GWAS identifies context-dependent QTL

In the accession panel, several QTL were detected for traits of interest. A major locus on chromosome 7 was associated with both growth habit and leaflet size, explaining more than 27% of the variance for each trait. This SNP was located near MS.gene036920, encoding a RING-type zinc finger protein. Homologues in *Arabidopsis thaliana* (L.) Heynh. and rice (e.g. AtRZFP34, OsRZFP34) regulate stomatal aperture and ABA responses (Hsu et al. 2014), suggesting a potential role in morphogenesis and water-use efficiency. QTL for growth habit are generally associated with related architectural traits, for example F. Zhang et al. (2019) identified several QTL for shoot diameter and branching on linkage group 7, traits potentially related to plant architecture. For leaflet traits, prior studies have used detailed measurements (length, width, area) to dissect size variation. He et al. (2019) reported 13 QTL for leaflet area, while Jiang, Yang, et al. (2022) identified seven candidate genes through combined QTL mapping and RNA-seq. None of their candidate genes overlapped physically with the loci identified in this study, suggesting here a novel region of interest.

A QTL for plant structure was detected on chromosome 6 in the accession panel, explaining 15.6% of the phenotypic variance. The candidate gene, MS.gene031124, encodes a ubiquitin conjugation factor E4, involved in protein degradation and cell survival (Koegl et al. 1999). In Arabidopsis, E4 family proteins such as AtUAP2 participate in ABA-mediated stress responses (Park et al. 2024). Previous work by McCord et al.(2014) identified two QTL for lodging (≥14% of phenotypic variation each) in a backcross population. However, the loci did not overlap physically with those identified here, likely due to differences in mapping populations and marker resolution. For growth rate, a minor QTL (3.8% of phenotypic variation) was identified on chromosome 8, near MS.gene61257, which encodes a calcium-binding protein with an EF-hand domain. EF-hand proteins are widely known to mediate calcium signalling during growth reactivation following stress (Lewit-Bentley and Réty 2000; Grabarek 2006).

In the Syn1, several QTL were detected for plant height, distributed across chromosomes 1, 2, 3, 4, 5, and 8, jointly explaining of 60% of phenotypic variance. The Syn1 also displayed longer linkage disequilibrium, as expected for an early-generation synthetic population with limited recombination (Li et al. 2014). While this feature does not explain the reduced phenotypic correlations, it is advantageous for GWAS because extended LD facilitates the detection of QTL even with a moderate marker density, as observed in this study. The locus with the strongest effect was located on chromosome 8 within MS.gene000456, a ubiquitin-associated domain protein, which may influence hormonal or structural growth pathways. Other candidate genes were implicated in signalling, chromatin organisation, and defence-related processes. Previous studies using biparental populations reported height QTL across all linkage groups, often co-localising with yield loci (Zhang et al. 2019; He et al. 2020; Jiang et al. 2022b). For leaflet size, QTL were identified on chromosomes 1, 2, and 4, jointly explaining 31% of the variance. Two QTL were not located close to genes but the QTL on chromosome 4 was located near MS.gene91452, though no functional annotation was available. Prior leaflet-related QTL studies, including the QTL detected in the accession panel, did not report overlapping loci with these regions (Jiang, Yang, et al., 2022). Unlike in the accession panel, where leaflet size and growth habit co-localised to a major QTL on chromosome 7, no overlapping QTL were detected for these two traits in the Syn1. This decoupling supports that recombination among divergent parents disrupted historical associations between traits.

Overall, QTL sets in the accession panel and the Syn1 did not overlap, underlining the strong dependence of trait–marker associations on genetic background, allele frequencies, and population history. The lack of overlap also reflects differences in phenotypic variance, the Syn1 displayed reduced variability for several traits, while in the accession panel, correction for strong population structure could have limited the detection of associations. Marker density (>100,000 SNP) provided informative resolution for both populations but cannot rule out the presence of undetected loci.

### 4.3. Genomic prediction accuracy depends on genetic relatedness and trait heritabilit

Genomic prediction varied substantially across the four prediction scenarios. Within the accession panel, prediction was highly effective for several traits. Growth habit and leaflet size achieved prediction abilities exceeding 0.8, even with small training subsets, while plant height, stem diameter, and plant structure reached moderate to high prediction abilities (0.5-0.7). These findings are consistent with previous studies reporting high prediction ability for traits such as plant height (PA = 0.65; (Jia et al. 2018)), autumn dormancy (PA > 0.75; in Pégard et al. (2023) and PA > 0.64 in F. Zhang et al. (2023)). For spring regrowth rate, prediction ability was weak (PA ≈ 0.30), which can be explained by its low heritability and polygenic nature, as also observed by Jia et al.(2018).

Within the Syn1 population, prediction ability was generally lower than in the accession panel but improved with increasing training set size. The highest performance was achieved for stem diameter and growth habit (PA = 0.37), while dormancy showed near-zero predictability. This reduced accuracy is likely attributable to both the limited genetic variability within the Syn1 and its small population size. Although the Syn1 was derived from three different accessions, recombination remains incomplete, and genetic diversity is narrower than in the accession panel. These factors restrict the model capacity to estimate reliable marker effects, a known limitation in early-generation breeding populations (H. Wang et al., 2024). A larger or more recombined synthetic population (e.g. Syn2) would likely improve performance.

Across-population predictions gave prediction abilities comparable to those obtained within the Syn1, generally below 0.30. Using a large and diverse training set to predict a narrow synthetic population did not improve accuracy but also did not perform worse than within-population prediction. This indicates that accession panels can provide predictive information across populations, although accuracy remains limited by the small size and reduced variance of the Syn1. This suggests that a single, large-based panel is valuable to predict traits in breeding pools whose genetic diversity is narrower than that of the panel but still included.

## 5. Conclusion

This study investigated the genetic determinants of lucerne traits relevant for ideotype breeding in living mulch systems, using both a diverse accession panel and a synthetic population. The accession panel captured broad phenotypic and genetic diversity structured by subspecies and cultivation status, while the Syn1 provided a complementary, less structured population where recombination generated novel trait combinations. Correlation patterns reflected these differences: strong associations in the accession panel illustrated historical selection and the maintenance of broad diversity, whereas reduced correlations in the Syn1 highlighted the potential to decouple traits through recombination. QTL identified through population-specific GWAS confirmed the polygenic nature of key architectural traits and their dependence on genetic background. Genomic prediction was highly effective in the accession panel, with accuracies above 0.80 for several traits, and moderately promising in the Syn1. Across-population predictions were comparable to those within the Syn1, showing that accession panels can inform prediction across populations, though accuracy remained constrained by the Syn1 limited size and variance. Overall, prediction ability depended not only on trait heritability but also on the diversity and relatedness between training and test sets

From a breeding perspective, these findings provide guidance for the development of lucerne ideotypes adapted to living mulch systems. Traits such as growth habit, leaflet size, and plant structure can be efficiently improved through genomic prediction within well-characterised and sufficiently large populations, while QTL may assist in selection within specific contexts. Synthetic populations underline the opportunity to recombine and decouple traits. This study provides a solid framework for breeding ideotypes that combine traits dedicated to living mulch use, i.e. reduced vigour with sufficient upright plant structure and adequate autumn dormancy.

## Acknowledgements

We thank the technical staff of the FERLUS and URP3F units at INRAE in Lusignan for their essential assistance in trial and laboratory monitoring.

## Statements and Declarations

The authors have no competing interests.

## Author contributions

B.J. and G.L. conceptualisation; Z.E.G., M.P., and B.J. methodology; Z.E.G., M.P., B.J., M.G., and I.A.-R. validation; Z.E.G. formal analysis; Z.E.G., F.S., and B.J. investigation; Z.E.G. writing-original draft preparation; Z.E.G., M.P., G.L., F.S., I.A.-R., and B.J. writing-review and editing; B.J., G.L., and M.P. supervision; B.J. and G.L. funding acquisition. All authors have read and approved the final manuscript.

## Data availability

The genotyping and phenotyping data generated and analysed during the current study will be available at https://doi.org/10.57745/SB0I2H when the paper is accepted.

## Supplementary materials

**Table S1.**
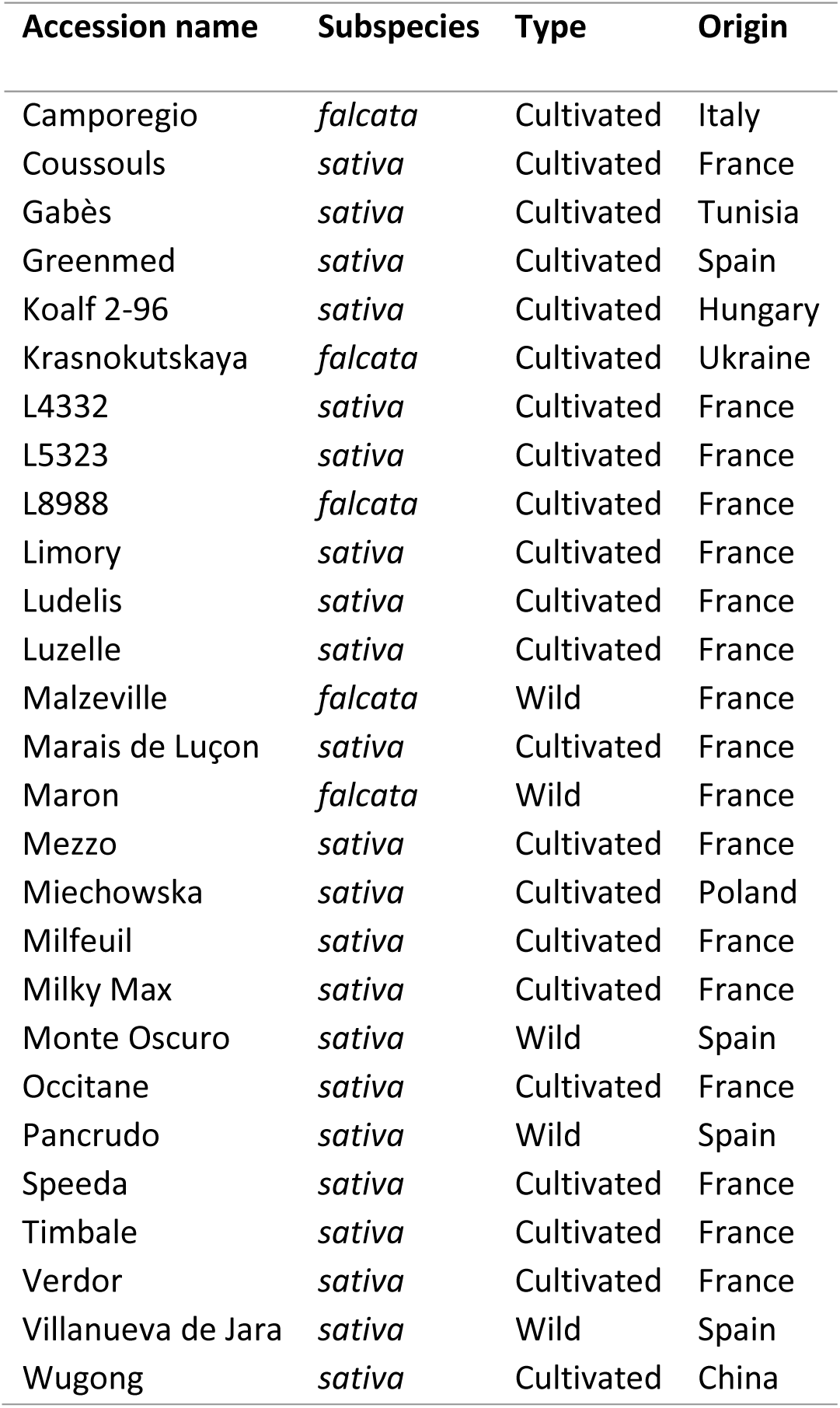
Characteristics of the 27 lucerne accessions, including subspecies, type (wild or cultivated), and geographical origin.

**Figure S1.**
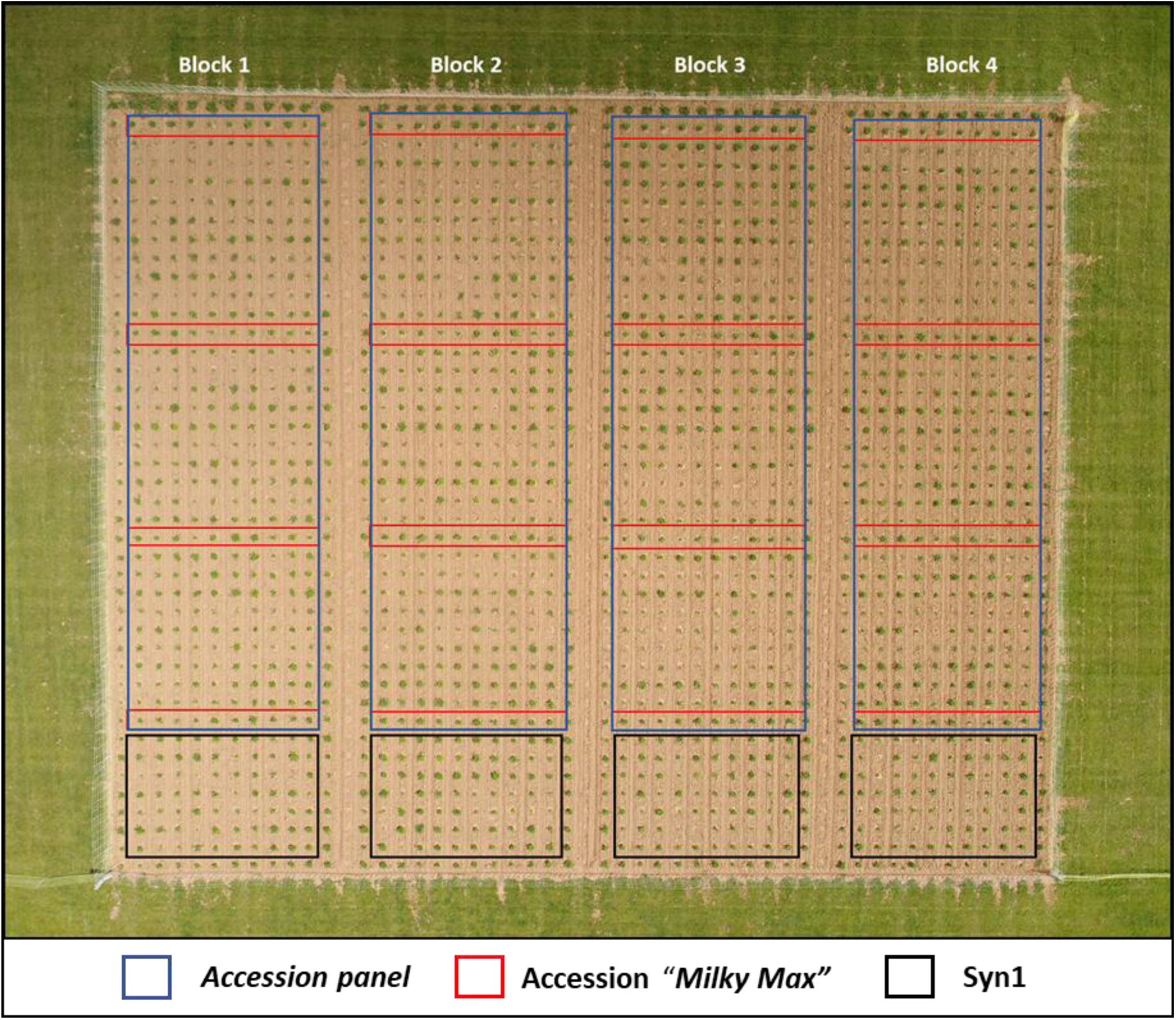
Field layout showing the four blocks, each containing 10 plants per accession; The “Milky Max” accession was repeated four times per block to assess spatial variability. The Syn1 population was planted at the end of each block. following the accession rows. All remaining plants surrounding the experimental plots served as border plants.

**Table S2.**
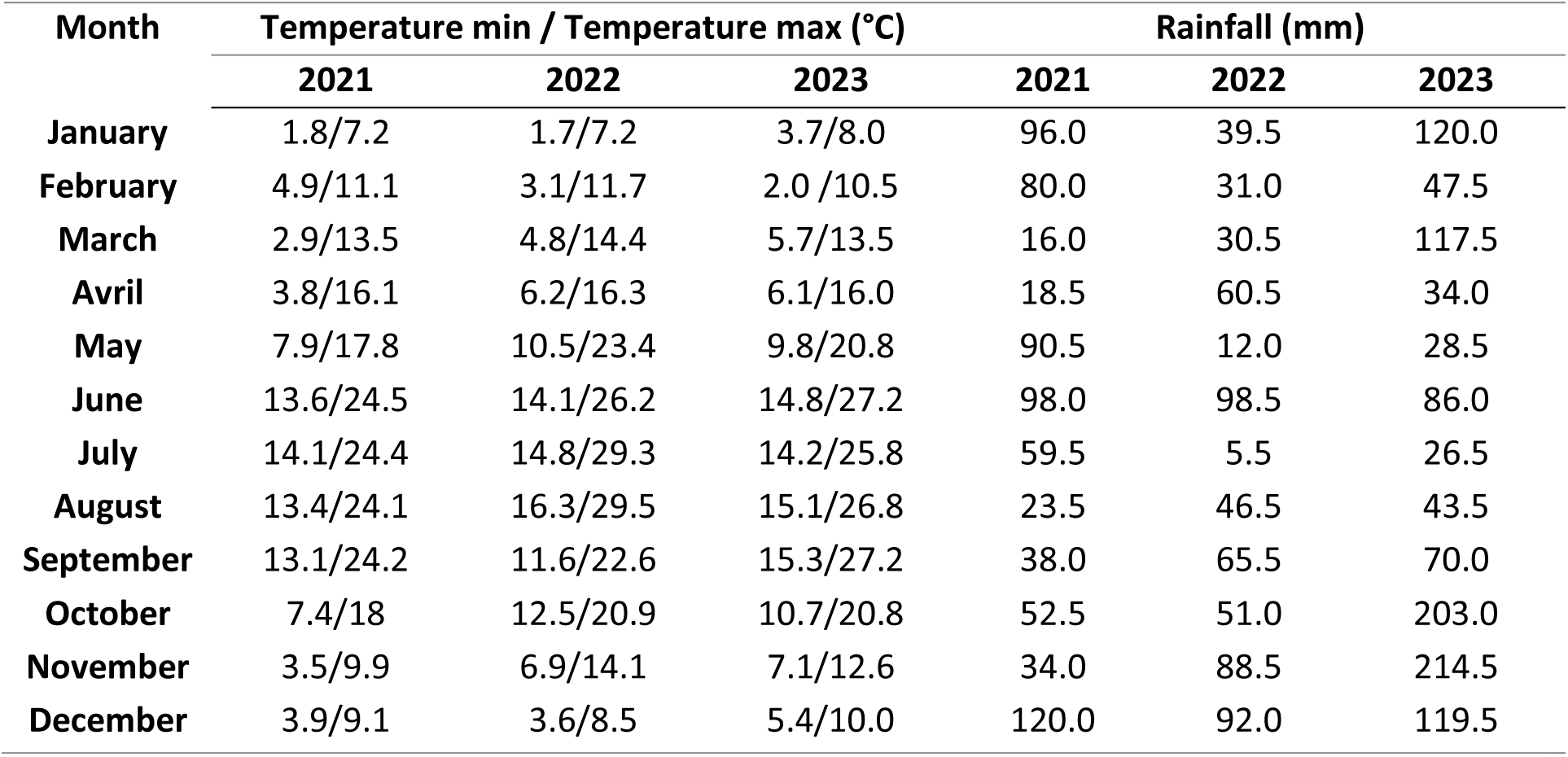
Climate data from April 2021 to June 2023.

**Figure S2.**
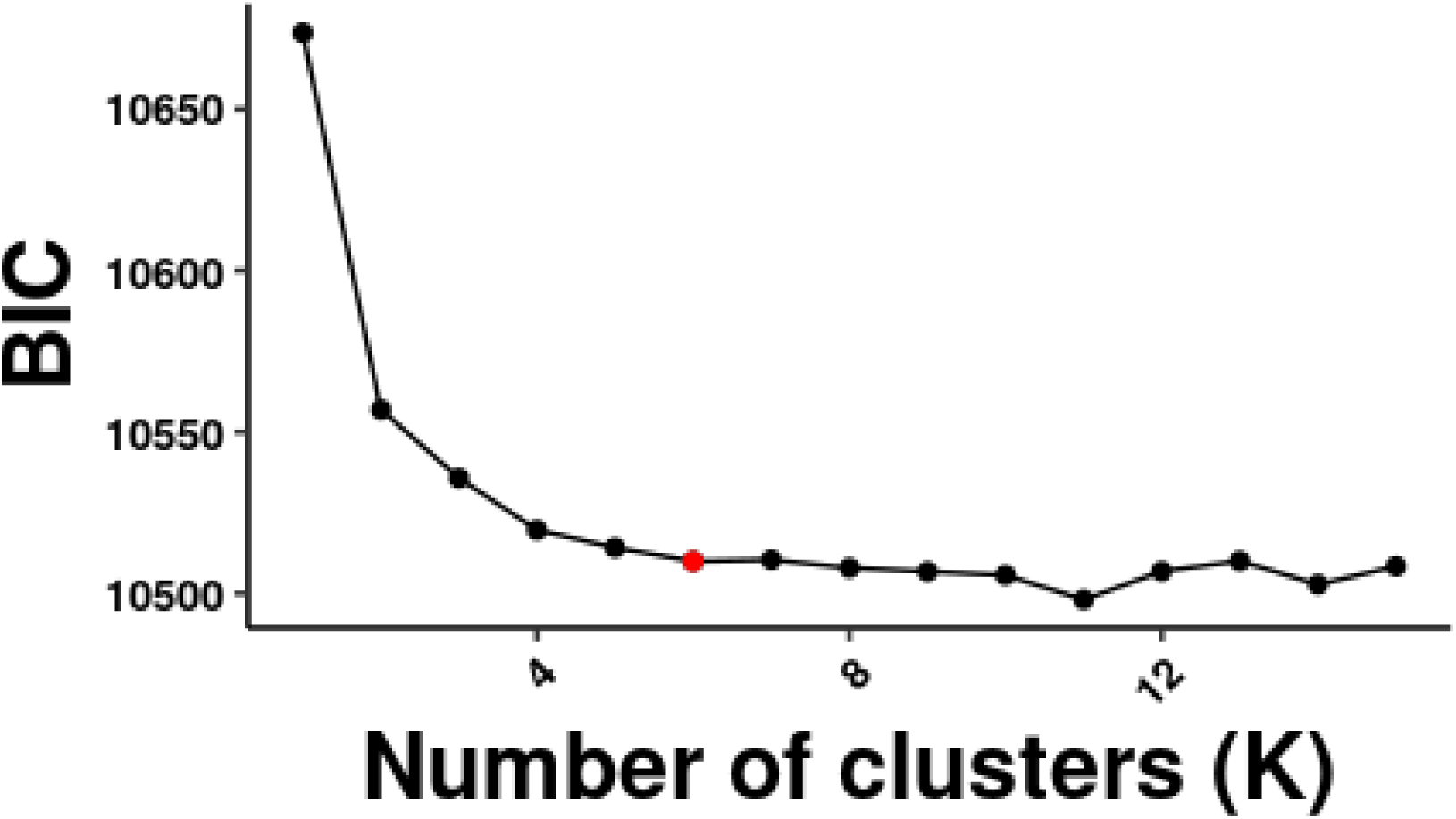
Bayesian Information Criterion (BIC) values for different numbers of clusters (K).

**Table S3.**
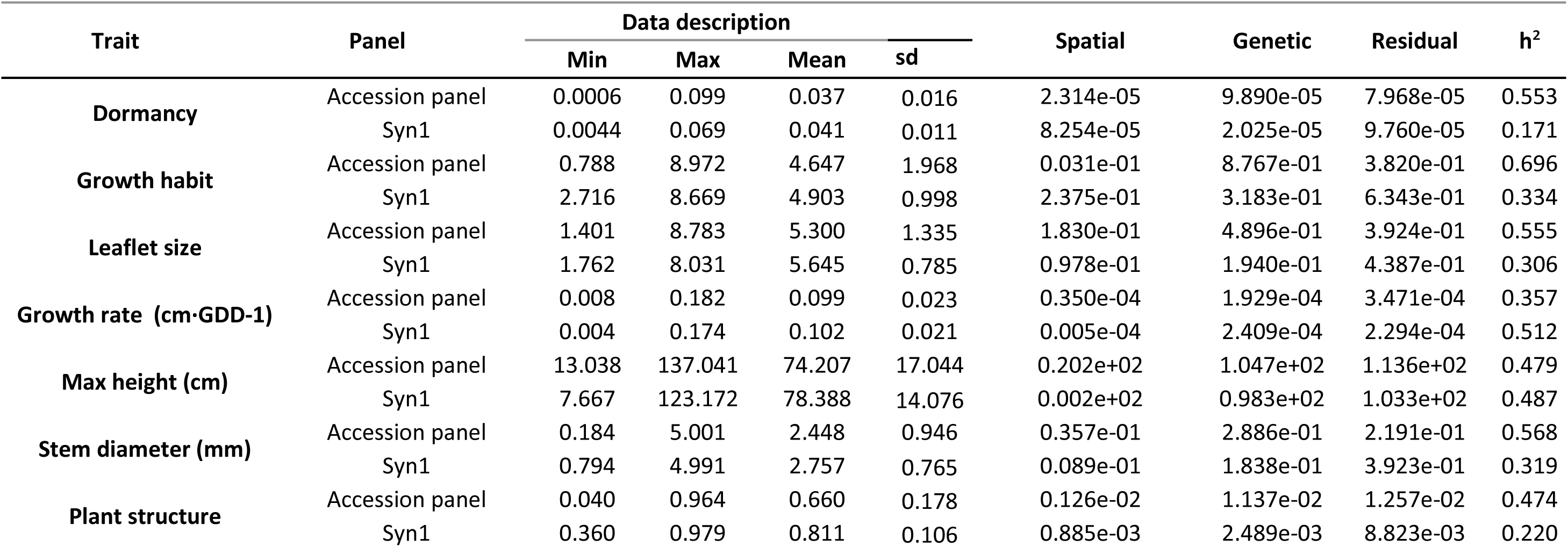
Summary of descriptive statistics (minimum. maximum. and mean) and heritability (*h*²) estimates for studied traits measured in the accessions panel and the Syn1 population. Heritability estimates were derived using the model described in Eq.1. Variance components (spatial. genetic. and residual) associated to these traits are provided.

**Figure 3S.**
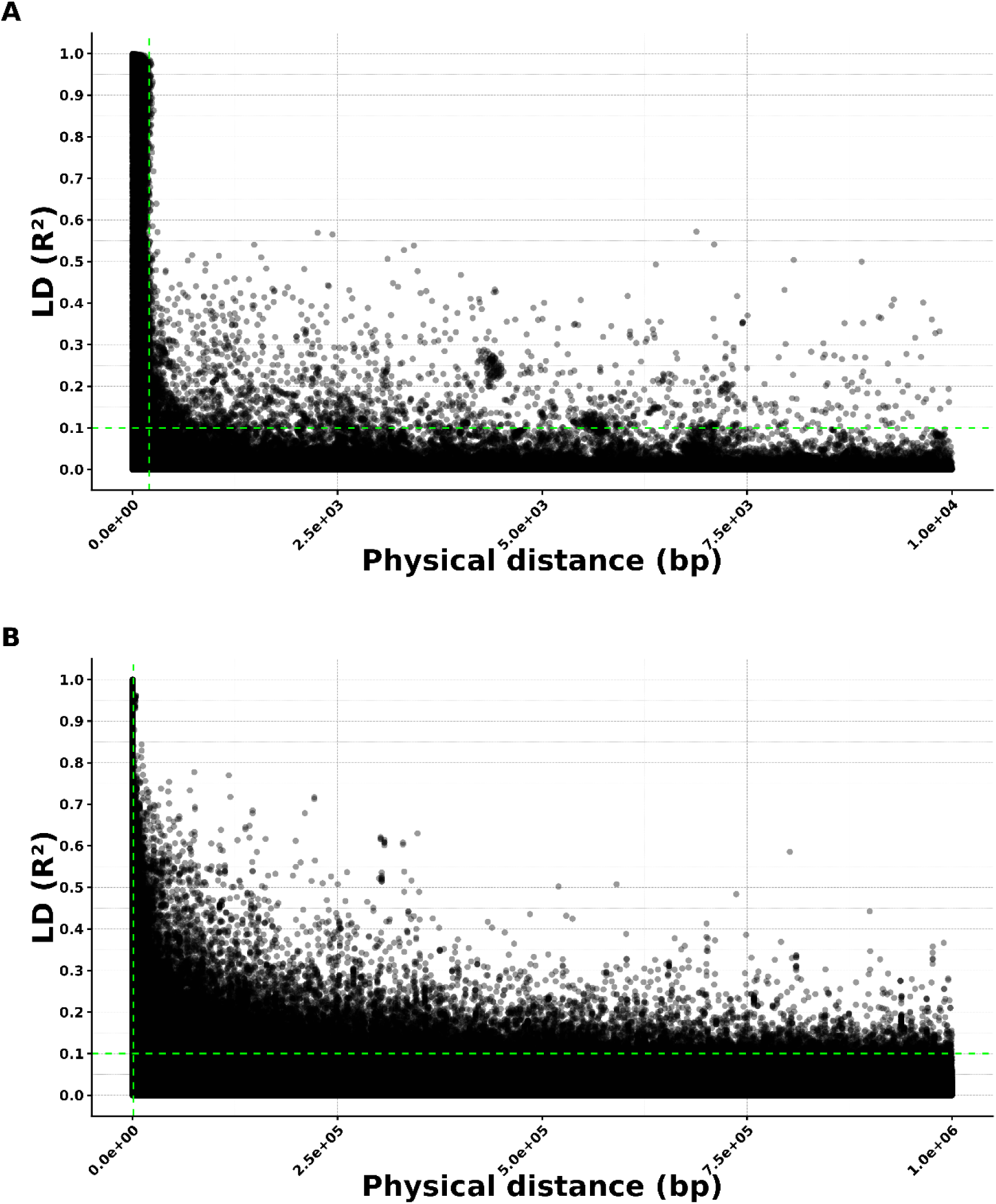
Linkage disequilibrium (LD) decay by chromosome. LD between SNP pairs is plotted against physical distance, with all chromosomes shown together in a single plot. The horizontal dashed line indicates the LD threshold (r² = 0.1), and the vertical dashed line marks 500 bp: (A) accession panel and (B) Syn1 population.

**Figure 4S.**
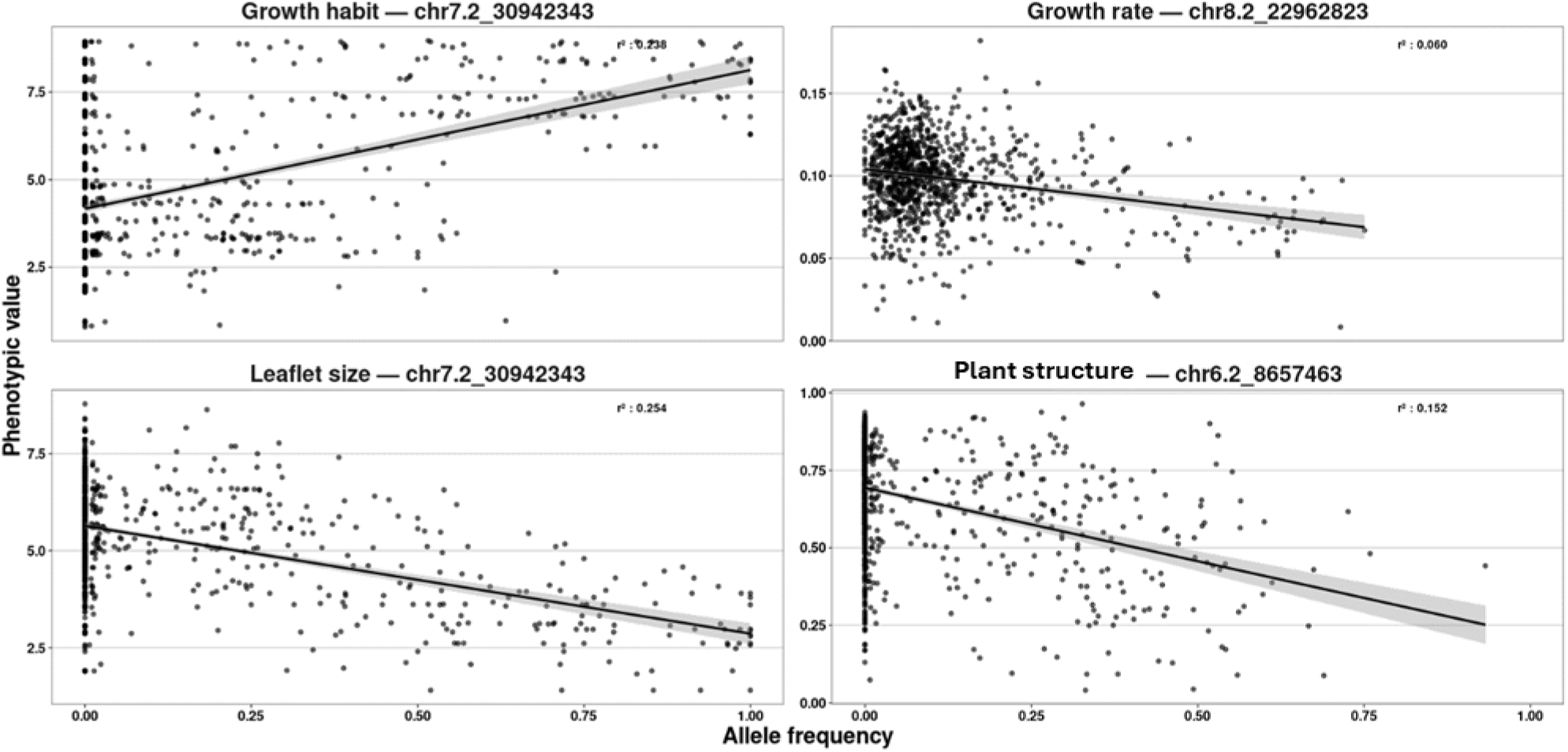
Allele frequency–phenotype associations for significant QTL identified in the accession panel.

**Figure 5S.**
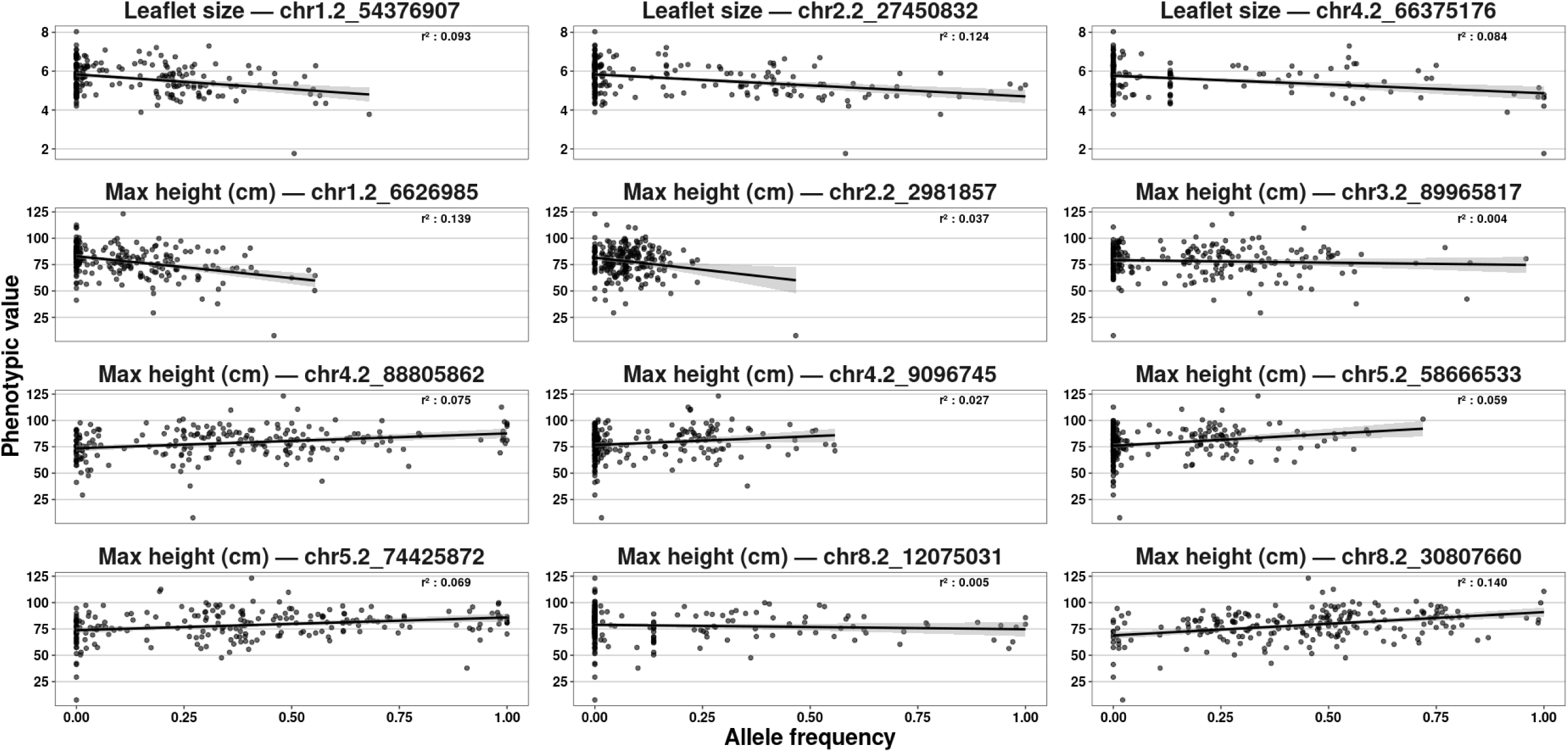
Allele frequency–phenotype associations for significant QTL identified in the Syn1 population

