## supplementary materials for "Genome-wide association study and genomic prediction of lucerne traits shaping living mulch performance"

Table S1 Characteristics of the 27 lucerne accessions, including subspecies, type (wild or cultivated), and geographical origin.

| Accession name | Subspecies | Type | Origin |
|---|---|---|---|
| Camporegio | *falcata* | Cultivated | Italy |
| Coussouls | *sativa* | Cultivated | France |
| Gabès | *sativa* | Cultivated | Tunisia |
| Greenmed | *sativa* | Cultivated | Spain |
| Koalf 2-96 | *sativa* | Cultivated | Hungary |
| Krasnokutskaya | *falcata* | Cultivated | Ukraine |
| L4332 | *sativa* | Cultivated | France |
| L5323 | *sativa* | Cultivated | France |
| L8988 | *falcata* | Cultivated | France |
| Limory | *sativa* | Cultivated | France |
| Ludelis | *sativa* | Cultivated | France |
| Luzelle | *sativa* | Cultivated | France |
| Malzeville | *falcata* | Wild | France |
| Marais de Luçon | *sativa* | Cultivated | France |
| Maron | *falcata* | Wild | France |
| Mezzo | *sativa* | Cultivated | France |
| Miechowska | *sativa* | Cultivated | Poland |
| Milfeuil | *sativa* | Cultivated | France |
| Milky Max | *sativa* | Cultivated | France |
| Monte Oscuro | *sativa* | Wild | Spain |
| Occitane | *sativa* | Cultivated | France |
| Pancrudo | *sativa* | Wild | Spain |
| Speeda | *sativa* | Cultivated | France |
| Timbale | *sativa* | Cultivated | France |
| Verdor | *sativa* | Cultivated | France |
| Villanueva de Jara | *sativa* | Wild | Spain |
| Wugong | *sativa* | Cultivated | China |

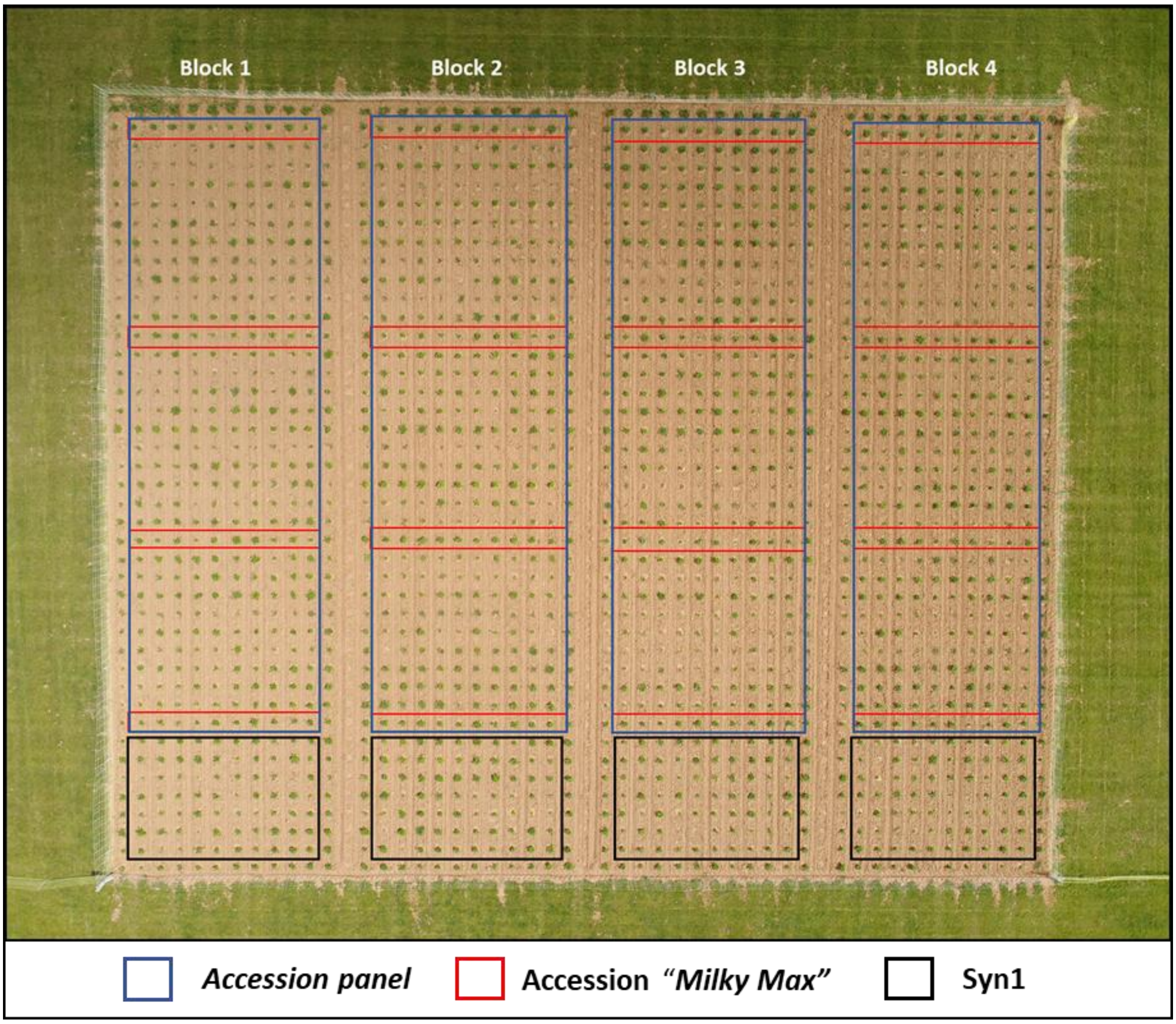


Figure S1 Field layout showing the four blocks, each containing 10 plants per accession; The "Milky Max" accession was repeated four times per block to assess spatial variability. The Syn1 population was planted at the end of each block. following the accession rows. All remaining plants surrounding the experimental plots served as border plants.

Table S2 Climate data from April 2021 to June 2023.

| Month | Temperature min / Temperature max (°C) | | | Rainfall (mm) | | |
|---|---|---|---|---|---|---|
| | 2021 | 2022 | 2023 | 2021 | 2022 | 2023 |
| **January** | 1.8/7.2 | 1.7/7.2 | 3.7/8.0 | 96.0 | 39.5 | 120.0 |
| **February** | 4.9/11.1 | 3.1/11.7 | 2.0 /10.5 | 80.0 | 31.0 | 47.5 |
| **March** | 2.9/13.5 | 4.8/14.4 | 5.7/13.5 | 16.0 | 30.5 | 117.5 |
| **Avril** | 3.8/16.1 | 6.2/16.3 | 6.1/16.0 | 18.5 | 60.5 | 34.0 |
| **May** | 7.9/17.8 | 10.5/23.4 | 9.8/20.8 | 90.5 | 12.0 | 28.5 |
| **June** | 13.6/24.5 | 14.1/26.2 | 14.8/27.2 | 98.0 | 98.5 | 86.0 |
| **July** | 14.1/24.4 | 14.8/29.3 | 14.2/25.8 | 59.5 | 5.5 | 26.5 |
| **August** | 13.4/24.1 | 16.3/29.5 | 15.1/26.8 | 23.5 | 46.5 | 43.5 |
| **September** | 13.1/24.2 | 11.6/22.6 | 15.3/27.2 | 38.0 | 65.5 | 70.0 |
| **October** | 7.4/18 | 12.5/20.9 | 10.7/20.8 | 52.5 | 51.0 | 203.0 |
| **November** | 3.5/9.9 | 6.9/14.1 | 7.1/12.6 | 34.0 | 88.5 | 214.5 |
| **December** | 3.9/9.1 | 3.6/8.5 | 5.4/10.0 | 120.0 | 92.0 | 119.5 |

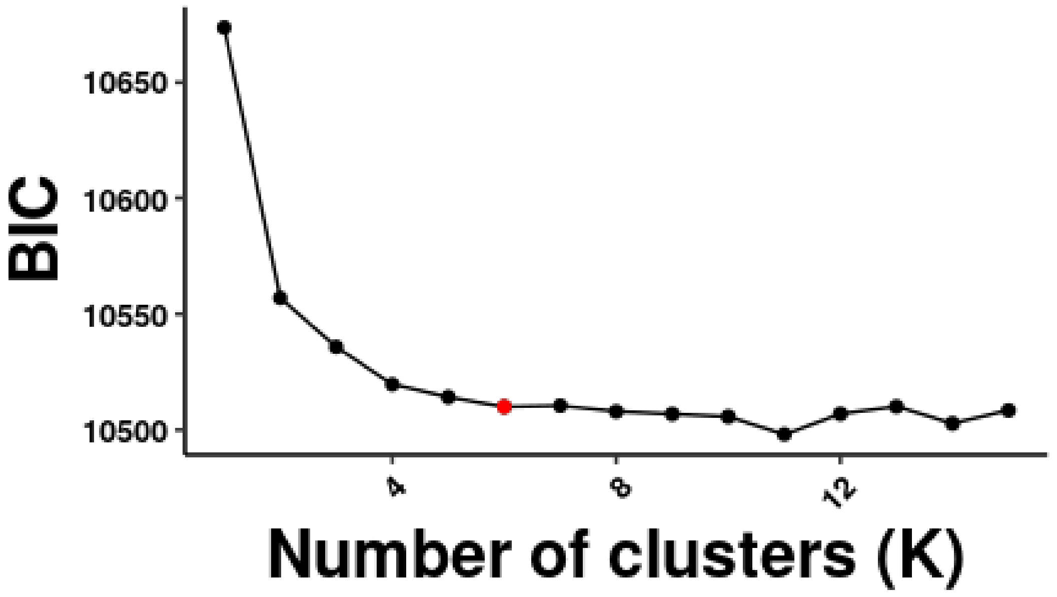


Figure S2 Bayesian Information Criterion (BIC) values for different numbers of clusters (K).

Table S3 Summary of descriptive statistics (minimum. maximum. and mean) and heritability ($h^2$) estimates for studied traits measured in the accessions panel and the Syn1 population. Heritability estimates were derived using the model described in Eq.1. Variance components (spatial. genetic. and residual) associated to these traits are provided .

| Trait | Panel | Data description | | | | Spatial | Genetic | Residual | $h^2$ |
|---|---|---|---|---|---|---|---|---|---|
| | | Min | Max | Mean | sd | | | | |
| **Dormancy** | Accession panel | 0.0006 | 0.099 | 0.037 | 0.016 | 2.314e-05 | 9.890e-05 | 7.968e-05 | 0.553 |
| | Syn1 | 0.0044 | 0.069 | 0.041 | 0.011 | 8.254e-05 | 2.025e-05 | 9.760e-05 | 0.171 |
| **Growth habit** | Accession panel | 0.788 | 8.972 | 4.647 | 1.968 | 0.031e-01 | 8.767e-01 | 3.820e-01 | 0.696 |
| | Syn1 | 2.716 | 8.669 | 4.903 | 0.998 | 2.375e-01 | 3.183e-01 | 6.343e-01 | 0.334 |
| **Leaflet size** | Accession panel | 1.401 | 8.783 | 5.300 | 1.335 | 1.830e-01 | 4.896e-01 | 3.924e-01 | 0.555 |
| | Syn1 | 1.762 | 8.031 | 5.645 | 0.785 | 0.978e-01 | 1.940e-01 | 4.387e-01 | 0.306 |
| **Growth rate (cm·GDD-1)** | Accession panel | 0.008 | 0.182 | 0.099 | 0.023 | 0.350e-04 | 1.929e-04 | 3.471e-04 | 0.357 |
| | Syn1 | 0.004 | 0.174 | 0.102 | 0.021 | 0.005e-04 | 2.409e-04 | 2.294e-04 | 0.512 |
| **Max height (cm)** | Accession panel | 13.038 | 137.041 | 74.207 | 17.044 | 0.202e+02 | 1.047e+02 | 1.136e+02 | 0.479 |
| | Syn1 | 7.667 | 123.172 | 78.388 | 14.076 | 0.002e+02 | 0.983e+02 | 1.033e+02 | 0.487 |
| **Stem diameter (mm)** | Accession panel | 0.184 | 5.001 | 2.448 | 0.946 | 0.357e-01 | 2.886e-01 | 2.191e-01 | 0.568 |
| | Syn1 | 0.794 | 4.991 | 2.757 | 0.765 | 0.089e-01 | 1.838e-01 | 3.923e-01 | 0.319 |
| **Lodging** | Accession panel | 0.040 | 0.964 | 0.660 | 0.178 | 0.126e-02 | 1.137e-02 | 1.257e-02 | 0.474 |
| | Syn1 | 0.360 | 0.979 | 0.811 | 0.106 | 0.885e-03 | 2.489e-03 | 8.823e-03 | 0.220 |

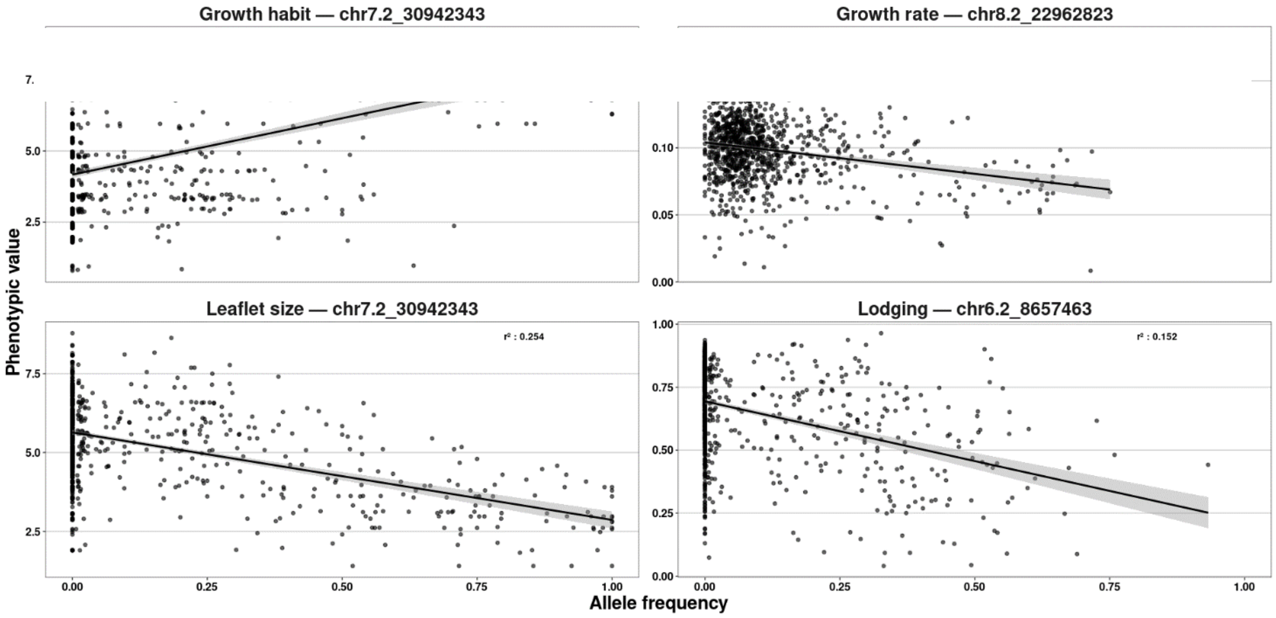


Figure 3S. Allele frequency–phenotype associations for significant QTL identified in the accession panel.

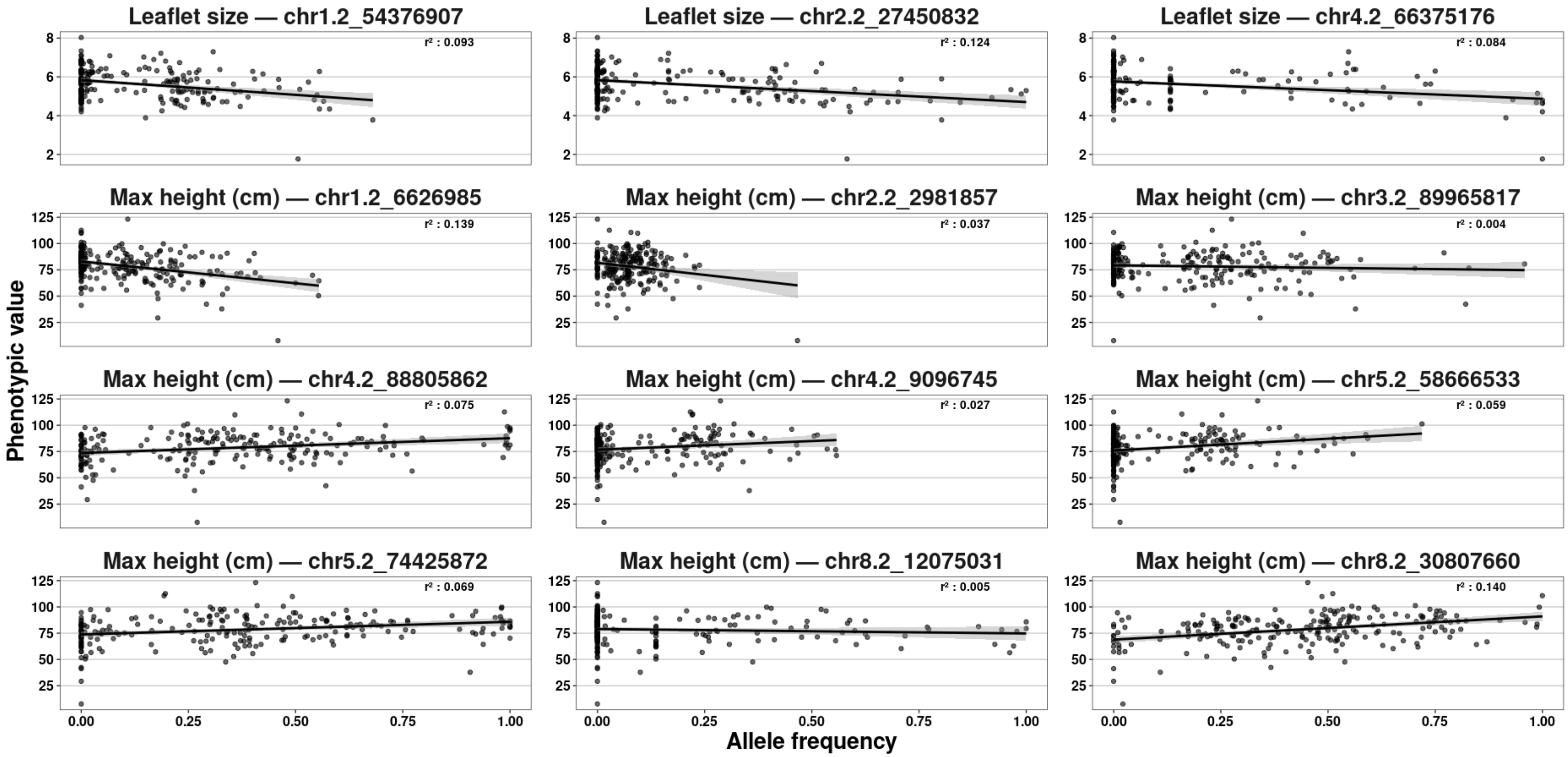


Figure 4S. Allele frequency–phenotype associations for significant QTL identified in the syn1 population.
